# Base-and-Sugar Dual-Frame Flow Matching for RNA Co-Design

**DOI:** 10.64898/2026.08.18.745543

**Authors:** Junzhe Li, Lijian Peng, Yuhao Li, Yize Zhou, Hanqun Cao, Cheng Tan, Shengchao Liu

**Affiliations:** Wave Intelligence Lab; CSE, CUHK

## Abstract

Current frame-based RNA generative models represent each nucleotide with a single rigid coordinate frame. However, our analysis of seven nucleotide frame constructions across 11,497 static RNA chains and 31,432 multi-state relation groups reveals a clear trade-off: base-centered frames better preserve base-mediated relations, whereas sugar-centered frames better support sugar-phosphate backbone reconstruction. Motivated by this complementarity, we introduce **DuetRNA**, a joint sequence-structure generation model for RNA co-design that performs SE(3) flow matching over coupled base-and-sugar frames. Empirically, we evaluate DuetRNA under both the inverse-folded (IF) backbone-design protocol and generated-sequence (GS) co-design protocol. DuetRNA improves scTM-validity over the strongest matched baselines by 7.67 and 4.33 % under IF and GS, respectively. Controlled ablations further support the necessity of the dual-frame modeling in DuetRNA. The official DuetRNA repository is https://github.com/XjunLi/DuetRNA.

## 1 Introduction

RNA molecules perform catalysis, molecular recognition, regulation, and assembly through structured three-dimensional conformations [Saenger, 1984, Garst et al., 2011, Vicens and Kieft, 2022, Ganser et al., 2019], making their structural design crucial to RNA therapeutics, synthetic biology, and programmable nanostructures [Damase et al., 2021, Han et al., 2017, Yesselman et al., 2019]. Recent generative models targeting this goal follow two related formulations. Structured-only design methods such as RNA-FrameFlow generate three-dimensional RNA backbones, with compatible sequences subsequently assigned through inverse folding [Anand et al., 2025]. RNA co-design instead generates sequence and structure simultaneously, modeling their compatibility within a single generative process [Rubin et al., 2025, Ma et al., 2025, Li et al., 2026b]. A key modeling choice in these generative frameworks is how nucleotide geometry is represented.

In frame-based protein structure modeling, a standard approach is to represent each residue with a single backbone-centered rigid frame [Jumper et al., 2021, Yim et al., 2023b,a]. Frame-based RNA generative models have largely inherited this formulation, likewise representing each nucleotide with a single rigid frame [Anand et al., 2025, Ma et al., 2025, Tarafder and Bhattacharya, 2026]. However, directly adopting the single frame from amino acid modeling to nucleotide modeling does not adequately utilize the specific structural attributes of RNA: nucleobases mediate pairing and stacking, whereas the sugar-phosphate backbone carries substantially greater conformational flexibility [Leontis and Westhof, 2001, Nissen et al., 2001, Parlea et al., 2016, Murray et al., 2003, Richardson et al., 2008, Clay et al., 2017]. These inherent differences raise a critical question: *How can RNA-specific geometry be better exploited for frame-based RNA co-design?*

### Our contributions

We first examine this question by revisiting the properties of frames in RNA structures. Across 11,497 static RNA chains and 31,432 multi-state relation groups, we compare seven base- and sugar-anchored frame constructions using interaction stability and atom-reconstruction criteria. Base-anchored frames most clearly stabilize canonical pairing and stacking orientation, while noncanonical relations remain slightly more heterogeneous. Sugar-centered frames retain the local coordinates needed to realize the ribose and phosphodiester chain. No tested single-frame representation simultaneously captures both roles effectively. A complementary dual-frame reconstruction analysis further shows that separating base and sugar geometry substantially reduces atom reconstruction error relative to the tested single-frame representations.

Motivated by this observation, we introduce DuetRNA, a dual-frame SE(3) flow-matching model for RNA co-design through joint sequence-structure generation. Each nucleotide carries a base-centered frame for residue-level organization and a sugar-centered frame for torsion-conditioned backbone reconstruction. The model jointly evolves both frames from noise while predicting nucleotide identities and local torsions, with a relative-pose objective coupling the two frames within each residue. Together, these variables enable direct atom-level reconstruction of complete RNA sequence-structure samples.

To evaluate DuetRNA, we consider two complementary de novo generation settings. The inverse-folded protocol (IF) measures backbone designability by assigning candidate sequences to generated structures and independently forward-folding them, whereas the generated-sequence protocol (GS) directly folds the model-generated sequence to test sequence-structure compatibility without post-hoc inverse folding. When trained on RNAsolo, DuetRNA achieves 48.67% IF scTM-validity, exceeding the strongest matched backbone-design baseline by 7.67 percentage points, and 38.50% GS scTM-validity, exceeding the strongest matched co-design baseline by 4.33 percentage points. Controlled ablations support the contributions of both base-sugar factorization and relative-pose coupling, while direct coordinate analyses reveal reduced novel-fold coverage and a localized glycosidic-linkage weakness. Together with our frame representation analysis, these results support dual-frame factorization as an effective inductive bias for RNA generative modeling, with base-centered frames capturing nucleobase-mediated organization and sugar-centered frames preserving the geometry required for backbone reconstruction.

## 2 Related Work

### Classical RNA modeling

RNA tertiary-structure modeling has long relied on fragment assembly, motif libraries, coarse-grained sampling, and knowledge- or physics-based energy functions. FARNA/FARFAR2, SimRNA, RNAComposer, and 3dRNA represent complementary versions of this paradigm, ranging from all-atom fragment assembly to coarse-grained statistical sampling and template-based reconstruction [Das and Baker, 2007, Watkins et al., 2020, Boniecki et al., 2016, Popenda et al., 2012, Wang et al., 2019]. These systems often encode pairing, stacking, and local conformational preferences explicitly, but their performance depends on sampling budget, template coverage, and the quality of auxiliary restraints. RNA-Puzzles and the RNA 3D Motif Atlas have documented both the progress of these methods and the remaining difficulty of noncanonical interactions and novel folds [Cruz et al., 2012, Miao et al., 2020, Parlea et al., 2016].

### Sequence-conditioned RNA tertiary-structure prediction

Deep learning has also transformed RNA tertiary-structure modeling, spanning learned structure scoring and sequence-conditioned prediction. ARES learns an RNA-specific geometric scoring function; DeepFoldRNA and trRosettaRNA predict geometric restraints; DRfold and DRfold2 combine residue frames, learned potentials, or denoising modules; RhoFold+ incorporates RNA language-model representations; and NuFold uses a flexible nucleobase-centered parameterization with explicit torsional structure [Townshend et al., 2021, Pearce et al., 2022, Wang et al., 2023, Li et al., 2023, 2026c, Shen et al., 2024, Kagaya et al., 2025]. RoseTTAFoldNA and AlphaFold 3 broaden this line to protein–nucleic-acid complexes and general biomolecular interactions [Baek et al., 2024, Abramson et al., 2024]. These models are valuable forward-folding oracles for design, but they do not by themselves define an unconditional distribution over RNA structures.

From a representation perspective, these predictors also reveal the importance of RNA-specific geometric parameterizations. DRfold learns nucleotide-wise local frames and inter-nucleotide geometric restraints, but uses a coarse-grained representation based on *P, C*4′, and glycosidic *N* atoms, with full-atom coordinates recovered afterward [Li et al., 2023]. NuFold is particularly relevant to our motivation: it adopts a flexible nucleobase-centered representation with explicit torsional modeling, and shows improved local geometry as well as accurate reconstruction of both *C*3′-endo and *C*2′-endo sugar puckers [Kagaya et al., 2025]. RNAbpFlow further demonstrates that flow matching can generate all-atom RNA structures under sequence and base-pair conditioning, while retaining a single nucleotide-level rigid-frame parameterization [Tarafder and Bhattacharya, 2026]. These results suggest that base orientation and sugar-backbone conformation play distinct geometric roles in RNA modeling. At the same time, systematic benchmarks show that current prediction methods still struggle on orphan RNAs and non-Watson-Crick interactions, with local interaction fidelity remaining a major bottleneck [Bahai et al., 2024]. This limitation is especially relevant for generation, where the model must construct plausible global folds together with fine-grained base-mediated interactions.

### Structure-first generation and inverse design

Structure-first design samples a geometric scaffold before assigning a compatible sequence. RNA-FrameFlow introduces SE(3) flow matching for *de novo* RNA backbone generation by representing each nucleotide as a rigid-body frame and predicting the remaining backbone atoms through torsional degrees of freedom [Anand et al., 2025]. Its established evaluation protocol further frames generation as a design problem: sampled backbones are passed through inverse folding with gRNAde and then forward folded with RhoFold to measure structural self-consistency [Anand et al., 2025, Joshi et al., 2025, Shen et al., 2022]. RFDpoly extends structure-first generation to RNA, DNA, proteins, and mixed biopolymer assemblies [Favor et al., 2025]. This two-stage organization is modular and can exploit a strong inverse-folding model, but sequence compatibility is introduced after the structural sample has already been generated. Moreover, their generative states remain organized primarily around backbone or residue-level geometry, leaving base organization to be recovered indirectly.

### RNA co-design and joint sequence–structure generation

RNA co-design aims to generate compatible RNA sequences and structures jointly rather than assigning sequence only after structure generation. MMDiff combines continuous geometric diffusion with discrete sequence diffusion for nucleic-acid and protein complexes, providing an early general framework for joint sequence-structure generation [Morehead et al., 2023]. RNAFlow uses inverse-folding-based flow matching for protein-conditioned RNA design [Nori and Jin, 2024]. RiboGen directly models RNA sequence and all-atom geometry through continuous and discrete flows, while RiboFlow combines residue frames, torsional variables, and sequence features for conditional RNA co-design [Rubin et al., 2025, Ma et al., 2025]. RiboDiff incorporates pre-trained structural priors into a joint sequence-structure diffusion model [Li et al., 2026b]. These works demonstrate that RNA co-generation is feasible. DuetRNA addresses a complementary representation question: whether one residue-level rigid frame can simultaneously organize nucleobase-mediated tertiary relations and support atomistic sugar-backbone realization.

### Geometric generative modeling

Continuous normalizing flows define generative dynamics through neural ordinary differential equations [Chen et al., 2018]; flow matching provides simulation-free vector-field regression, and Riemannian flow matching extends the construction to manifolds such as SO(3) and SE(3) [Lipman et al., 2023, Chen and Lipman, 2024]. Geometric neural networks preserve rigid-motion symmetries through equivariant attention or message passing, including SE(3)-Transformers, EGNNs, geometric vector perceptrons, and PaiNN [Fuchs et al., 2020, Satorras et al., 2021, Jing et al., 2021, Schütt et al., 2021]. Equivariant diffusion has been applied to atomic molecules, while residue-frame diffusion and flow matching have driven protein backbone generation in SE(3) Diffusion, FrameFlow, Genie, and RFdiffusion [Hoogeboom et al., 2022, Yim et al., 2023b,a, Lin and AlQuraishi, 2023, Watson et al., 2023]. DuetRNA follows this rigid-frame tradition but uses a product state with separate base and sugar objects, building on the role of frames in AlphaFold-style invariant point attention [Jumper et al., 2021].

### RNA structural data and validation

RNAsolo provides cleaned PDB-derived RNA structures, whereas RNA3DB constructs sequence- and structure-aware splits for machine-learning evaluation [Adamczyk et al., 2022, Szikszai et al., 2024]. Global comparison and clustering commonly rely on US-align and related TM-score procedures [Zhang et al., 2022]. Folding-based self-consistency can use RNA-specific predictors such as RhoFold+ or broader biomolecular predictors such as Boltz-1 [Shen et al., 2024, Wohlwend et al., 2024], but these scores do not establish atomistic stereochemical validity. MolProbity clash analysis, RNA backbone conformers, sugar-pucker criteria, and canonical bond geometry provide complementary local checks [Chen et al., 2010, Williams et al., 2018, Richardson et al., 2008, Clay et al., 2017, Gelbin et al., 1996]. DSSR and FR3D provide established interaction annotations and local geometric conventions for base pairs, stacking, and recurrent motifs [Lu et al., 2015, Sarver et al., 2008, Leontis and Westhof, 2001]. Our evaluation therefore separates folding-based designability, distributional coverage, and direct coordinate-level geometry.

## 3 Preliminaries

### 3.1 Rigid-Body Motions and SE(3)

#### The Rotation Group

SO(3). A rotation of a rigid body is represented by a 3 × 3 matrix *R* that preserves distances and orientations. All such matrices form the *special orthogonal group*

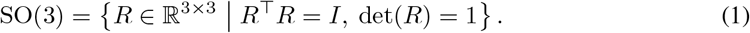

SO(3) is a three-dimensional *Lie group* and can be viewed as a Riemannian manifold once equipped with a suitable metric. Since SO(3) is not a Euclidean vector space, operations such as interpolation, averaging, and regression must respect its manifold structure.

#### The Special Euclidean Group

SE(3). A full rigid-body pose adds a translation *x* ∈ ℝ^3^ to the orientation. The pair belongs to the *special Euclidean group* [Barfoot, 2017]

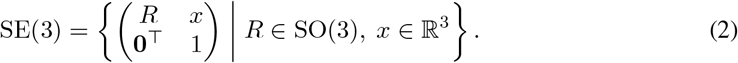

We write a pose as *T* = (*R, x*). Composition and inversion are

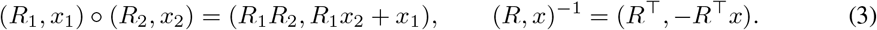

SE(3) is also a Lie group. Its tangent space at the identity, the Lie algebra *se*(3), is a six-dimensional vector space. An element of *se*(3) is parameterized by a six-dimensional coordinate *ξ* = (*ω, v*), with *ω* encoding the angular component and *v* the translational component. The exponential and logarithmic maps

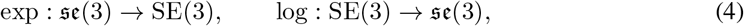

provide a local correspondence between the Lie algebra and the group. These maps enable local computations in the Lie algebra while representing configurations as points on SE(3).

### 3.2 Flow Matching on Riemannian Manifolds

Let ℳ be a Riemannian manifold and let *p*_data_ denote the unknown data distribution on ℳ. Flow matching [Lipman et al., 2023] learns a time-dependent vector field whose induced flow transports a simple prior distribution *p*_prior_ on ℳ to the data distribution through an ordinary differential equation [Chen et al., 2018].

#### Probability flow ODE

A time-dependent vector field *v*_*t*_ : ℳ → *T* ℳ defines an ordinary differential equation as

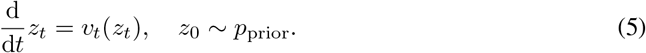

Here, *T*ℳ denotes the tangent bundle of ℳ, and *v*_*t*_(*z*) ∈ *T*_*z*_ ℳ for each *z* ∈ ℳ. Integrating this ODE from *t* = 0 to *t* = 1 yields a flow *ψ*_*t*_ such that *z*_*t*_ = *ψ*_*t*_(*z*_0_).

#### Interpolation construction

Since the marginal probability path *p*_*t*_ = (*ψ*_*t*_)_#_*p*_prior_ is intractable, we adopt a simulation-free training strategy. For each data point *z*_1_ ~*p*_data_, we draw a random noise sample *z*_0_ ~*p*_prior_ and construct an interpolation *z*_*t*_ between them. A natural choice on ℳ is the geodesic path [Chen and Lipman, 2024]:

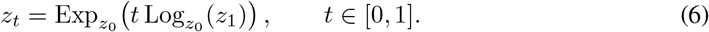

where Exp_*z*_ and Log_*z*_ denote the Riemannian exponential and logarithmic maps at point *z*.

#### Training via Endpoint parameterization

We adopt an *endpoint parameterization* [Lipman et al., 2023], in which the network predicts the terminal point 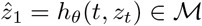 from interpolation point *z*_*t*_. This parameterization allows the velocity to be constructed geometrically through the logarithmic map:

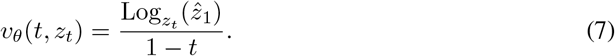

The learning objective is then written in the tangent space at *z*_*t*_ as

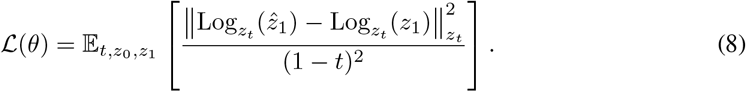

#### Inference

At inference time, source samples are integrated from *t* = 0 to *t* = 1 using the velocity reconstructed from the predicted endpoint.

An RNA chain with one rigid frame per residue lies on SE(3)^*N*^. DuetRNA assigns two frames to each residue, so its continuous state lies on the product manifold

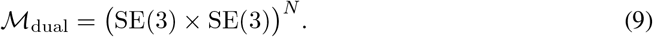

Translations are interpolated linearly in ℝ^3^, rotations follow geodesics on SO(3), and the two frame channels are coupled by a shared denoiser and a within-residue relative-pose objective described next in Section 4.

## 4 Method

We propose DuetRNA, a base-and-sugar dual-frame generative framework for RNA co-design. As shown in Figure 2, each nucleotide is described by two physically meaningful rigid objects: a nucleobase-centered frame for residue-level organization and a sugar-centered frame for sugar-backbone realization. The model learns a joint SE(3) flow over the two frames, predicts nucleotide identity and intra-residue torsional variables from coupled geometric features, and decodes the final state into an atom23 heavy-atom representation.

### Problem formulation

Let *C* = {A, U, G, C} denote the RNA nucleotide alphabet, *s* = (*s*_1_, …, *s*_*N*_) ∈ *C*^*N*^ the nucleotide sequence, and *ϕ* = (*ϕ*_1_, …, *ϕ*_*N*_) the backbone torsional variables. Our model jointly designs the structure and sequence of RNA by learning *p*_*θ*_(*T* ^base^, *T* ^sugar^, *s, ϕ*), with the factorization

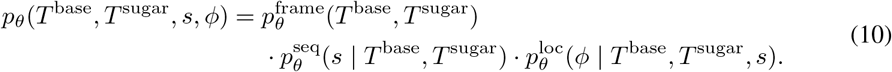

The following subsections follow the generative pipeline. Section 4.1 first transforms raw RNA data into a dual-frame representation. Section 4.2 estimates the joint distribution over base and sugar frames. Section 4.3 and Section 4.4 describe sequence prediction and intra-residue torsion estimation for atom23 completion. Section 4.5 defines the deterministic atom23 completion path, and Section 4.6 summarizes the training objective and model architecture.

### 4.1 Dual-Frame Representation of RNA

Rigid frames are widely used to compress molecular geometry for SE(3)-aware learning. AlphaFold2 and subsequent diffusion and flow-matching generative models operate on residue-level frames, while RigidSSL, AssembleFlow, InertialAR, and InertialGenome extend rigidity-aware or inertial-frame representations to protein pretraining, molecular assembly, whole-molecule canonicalization, and chromosome modeling, respectively [Jumper et al., 2021, Yim et al., 2023b,a, Anand et al., 2025, Ni et al., 2026, Guo et al., 2025, Li et al., 2026a, Zhou et al., 2026]. Together, these developments establish the frame definition as a modeling inductive bias rather than a neutral coordinate convention.

Our systematic analysis of residue-level coordinate frames (Appendix B) indicates that no single nucleotide frame simultaneously captures base-mediated inter-residue geometry and the local sugar-backbone geometry required for reconstruction. Base-centered frames are better aligned with many nucleobase-mediated relations, most clearly canonical pairs and stacking orientations, whereas sugar-centered frames retain the local coordinates needed for atomistic backbone reconstruction. We therefore represent residue *i* with a base frame 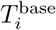 for base-mediated organization and a sugar frame 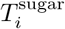 for torsion-conditioned sugar-backbone realization.

#### Deployed frame definitions

As shown in Figure 1, DuetRNA constructs one sugar frame and one base frame for each nucleotide. Specifically, the sugar frame *T* ^sugar^ uses **Sugar-GS**, a three-atom Gram-Schmidt frame anchored at *C*4′ and built from (*O*4′, *C*4′, *C*3′). The base frame *T* ^base^ uses **Base-Plane**, a chemically anchored nucleobase-plane frame whose origin is the glycosidic connection atom (*N* 9 for purines and *N* 1 for pyrimidines) and whose orientation is defined by the fitted base plane and an in-plane chemical axis. The exact coordinate definitions of these two deployed frames are given in Appendix B.

**Figure 1:**
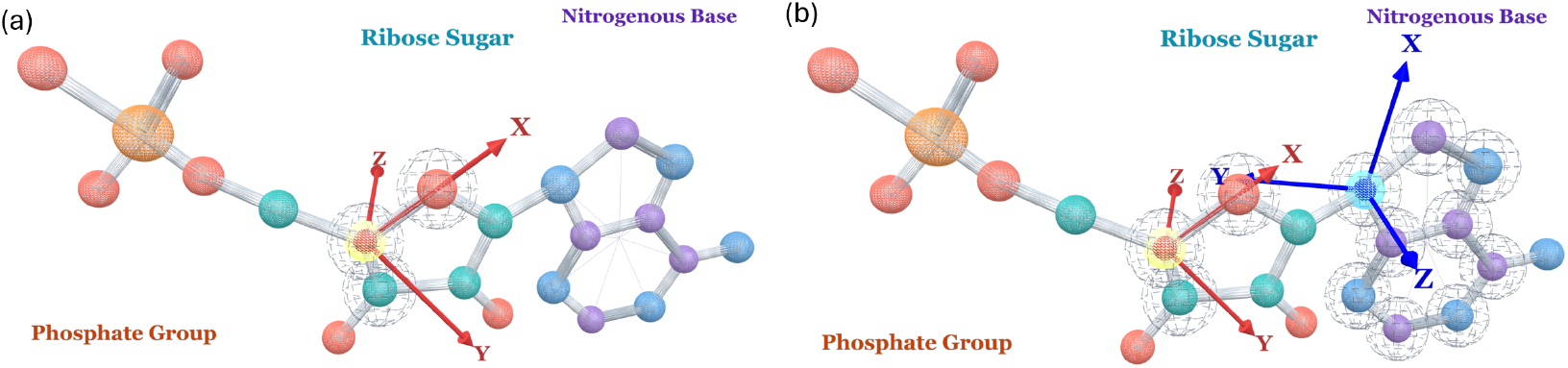
Comparison of RNA frame representations. (a) A single sugar Gram-Schmidt frame (Sugar-GS) is constructed from (*O*4′, *C*4′, *C*3′). (b) DuetRNA uses a Sugar-GS frame together with a chemically anchored Base-Plane frame.

**Figure 2:**
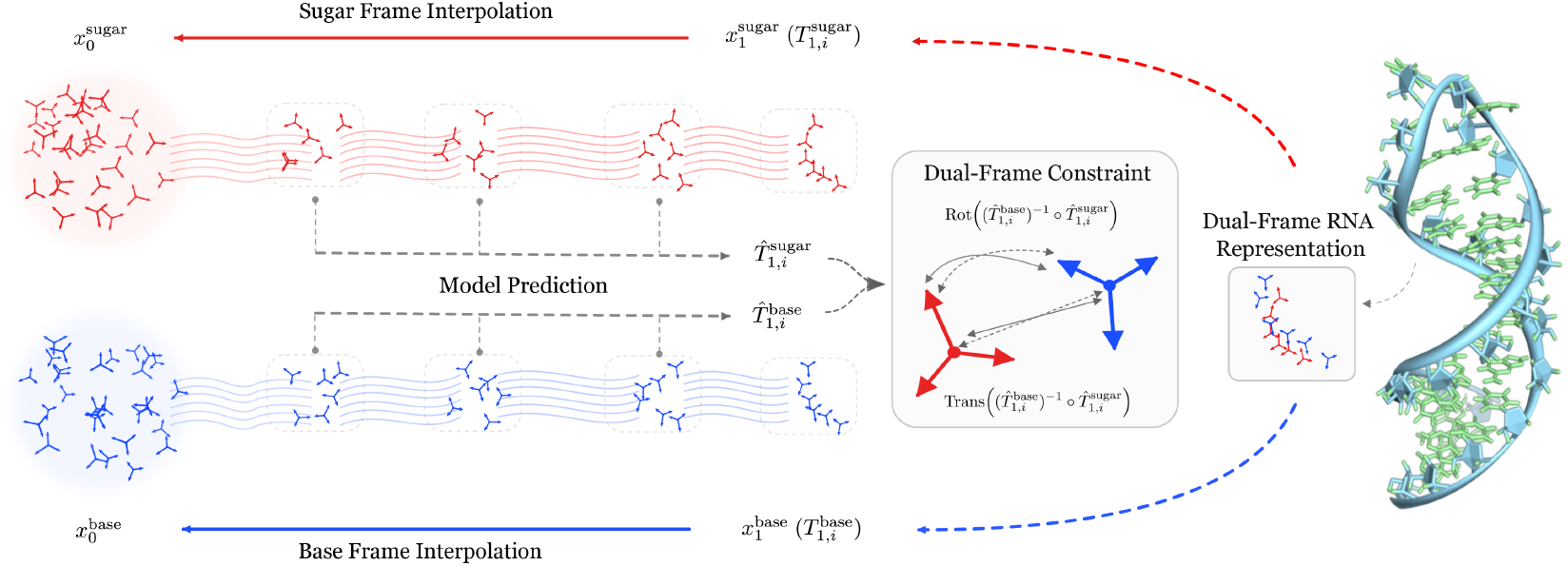
Overview of DuetRNA. In DuetRNA, each nucleotide is represented by two coupled SE(3) frames: a sugar frame and a base frame. The model jointly predicts terminal sugar and base frames from noisy interpolants, while a dual-frame constraint matches the predicted base-to-sugar relative translation and rotation to their ground-truth relative pose. This couples the two generated frames into a physically consistent dual-frame RNA representation for downstream sequence and atom-level structure reconstruction.

#### Dual-frame state

For a chain of length *N*, the RNA geometric state is thus described by

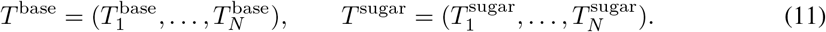

### 4.2 Dual-Frame Flow Matching

For an RNA molecule with *N* residues, the learning target for continuous geometry is the pair of frames

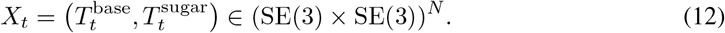

#### Source endpoints and dual-frame interpolation

For *f* ∈ {base, sugar}, we sample a source endpoint 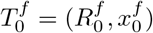 from an unconditional source distribution with Gaussian translations and uniform rotations on SO(3) as the prior distribution *p*_0_, and take a clean data frame 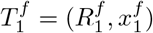 as the target distribution *p*_1_. We then define the interpolation path 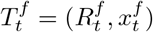 at time *t* as:

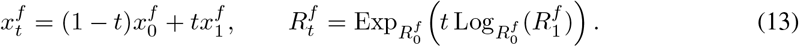

#### Reparameterized vector field estimation

Given 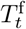 and time *t*, the model estimates the vector field by predicting terminal frame state 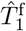. The implied translation and rotation vector fields from this reparameterization are

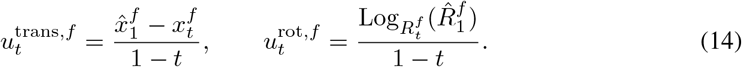

#### Learning objectives for vector field estimation

The estimation objective designed for Dual-Frame flow matching is

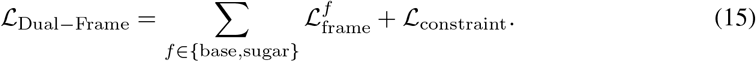

Here the rigid term is the combined SE(3) flow matching loss described in RNA-FrameFlow:

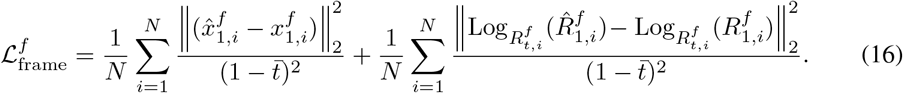

Here 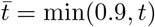 clips the flow time in the denominator. To ensure the two frames preserve the within-residue relationship during the flow matching process, we further introduce the reparameterized Dual-Frame constraint term in Equation (15) which supervises terminal relative translation and rotation at the predicted terminal state (*t* = 1).

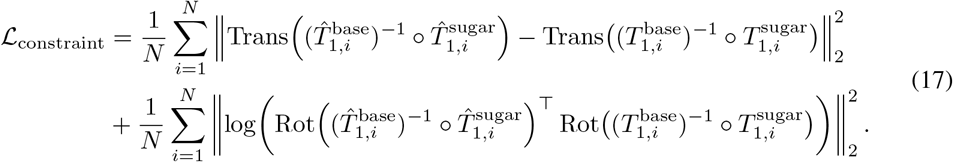

Here Trans(·) and Rot(·) extract the translation and rotation of an SE(3) transform, respectively.

### 4.3 Sequence Prediction

The sequence head estimates 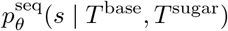 as a residue-wise categorical distribution. Let *π*_*i*_(*c*) be the predicted probability of nucleotide *c* ∈ {*A, U, G, C*} at residue *i*:

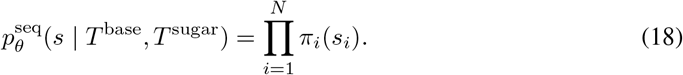

The maximum-likelihood objective is the residue-wise negative log-likelihood

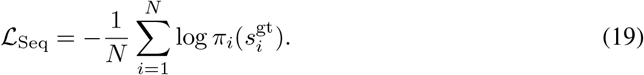

### 4.4 Intra-residue Torsion Estimation

The intra-residue torsional variables are the torsion angles required by atom23 completion:

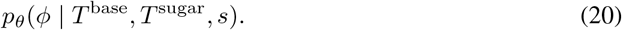

Let *K* = 8 be the number of torsion angles used by the completion map. These angles are predicted from the coupled geometric features by

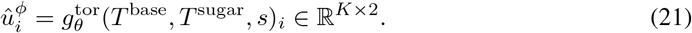

During training, these internal-coordinate variables are supervised both directly and through the decoded atom23 structure described in Section 4.5. The main local objective is

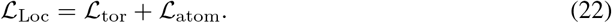

The torsion supervision uses a sine-cosine parameterization:

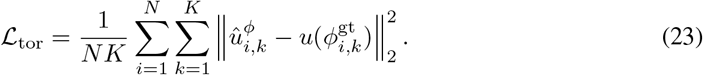

where *u*(*ϕ*_*i,k*_) = (sin *ϕ*_*i,k*_, cos *ϕ*_*i,k*_) ∈ *S*^1^.

The atom reconstruction loss is

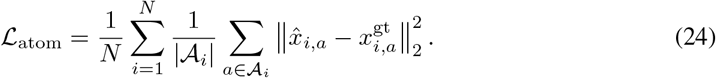

where *A*_*i*_ is the valid heavy-atom subset for residue *i* under the atom23 mask. Additional decoded-geometry supervision terms used in ablations are summarized in Appendix C.

### 4.5 Atom23 Completion

Atom23 denotes a fixed-width heavy-atom representation with up to 23 nucleotide-specific atom slots. Residue-specific absent atoms are masked, and hydrogens are excluded. Atom23 completion is a deterministic mapping from the predicted geometric variables to this compact heavy-atom representation:

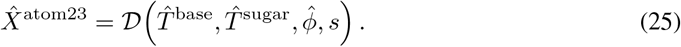

The sugar/backbone atoms are generated from the predicted sugar frame 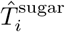 together with the predicted torsions (following RNA-FrameFlow [Anand et al., 2025]), whereas the base atoms are placed from canonical nucleotide-specific templates attached to the predicted base frame 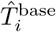. The final output is a compact atom23 heavy-atom representation rather than a hydrogen-complete all-atom model.

### 4.6 Training Objectives and Model Architecture

#### Objective

Overall, the training loss is the sum of three components from Sections 4.2 to 4.4:

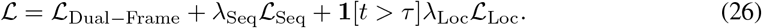

where ℒ_Dual-Frame_, ℒ_Seq_ and ℒ_Loc_ are described in Equations (15), (19) and (22), respectively. We use *τ* = 0.25 to gate ℒ_Loc_ to delay decoded-geometry supervision until the noised frames have moved beyond the earliest part of the interpolation path. The loss weights and auxiliary loss ablations are listed in Appendix C.

#### Architecture

The network architecture contains a coupled dual-frame geometric trunk and two output heads for sequence estimation and torsion angle estimation. The trunk takes the base and sugar frames as input, refines their coupled representations across depth, and exposes final features to the estimation heads described above. A more detailed discussion is provided in Appendix A.

## 5 Experiments

### 5.1 Experimental Setup Training data

The primary experiments use DuetRNA trained on the RNAsolo dataset [Adamczyk et al., 2022], following the same choice of RNA-FrameFlow [Anand et al., 2025]. This keeps the training-corpus alignment for the comparison with previous methods. As another version, we also train DuetRNA on RNA3DB [Szikszai et al., 2024], a PDB-derived RNA structure collection curated for deep-learning benchmarks with sequence- and structure-aware nonredundant splits. Results from the RNA3DB development run are retained for the auxiliary-objective ablation, sampler, and training-budget analyses in Appendices C.2–C.4.

#### Evaluation grids

The RNA-FrameFlow grid contains lengths 40 to 150 nt at intervals of 10, with 50 samples per length and 600 samples in total. The RiboFlow comparison uses lengths 50, 70, 90, 110, 130, and 150 nt, with 100 samples per length. RiboGen follows the same grid as RNA-FrameFlow. DuetRNA is rerun on each grid instead of comparing values obtained from different length distributions.

#### Sequence protocols

The inverse-folded-sequence protocol, denoted IF, assigns eight candidate sequences to each generated structure with gRNAde and folds them with RhoFold, following the RNA-FrameFlow evaluation protocol [Anand et al., 2025, Joshi et al., 2025, Shen et al., 2022]. All RhoFold-based evaluations use the same `RhoFold_pretrained.pt` checkpoint used in the RNA-FrameFlow evaluation pipeline. The best self-consistency score measures whether the generated backbone is designable by an external sequence model. The generated-sequence protocol, denoted GS, folds the sequence proposed by the generative model itself. It removes gRNAde and measures compatibility of the generated sequence-structure pair. We use Boltz-1 for the matched RiboGen comparison and report the RhoFold GS result separately [Wohlwend et al., 2024, Rubin et al., 2025].

#### Metrics and statistics

A sample is treated as valid when forward-folding self-consistency score scTM ≥ 0.45, following RNA-FrameFlow [Anand et al., 2025]. Diversity has three complementary definitions. Let *G* denote the set of all *N* generated structures and *V* ⊆ *G* the subset of *M* valid structures. We cluster structures with qTMclust using a TM-score cutoff of 0.45 [Zhang et al., 2022], and let *K*(*S*) denote the number of structural clusters obtained from a set *S*. We distinguish three quantities: all-sample diversity, *D*_all_ = *K*(*G*)*/N*; valid-set diversity, *D*_valid_ = *K*(*V*)*/M*; and unique-valid yield, *Y*_UV_ = *K*(*V*)*/N*. The first measures structural mode coverage across all generated samples, the second measures diversity conditional on designability, and the third measures the number of distinct designable modes obtained per generation attempt. Novelty is measured by the reference-set pdbTM. For each valid sample, we compute the US-align TM-score [Zhang et al., 2022] (RNA mode, *C*3′ representation, normalized by the length of training structures) against all training structures of the corresponding corpus, take the maximum over the training set, and average the resulting per-sample maxima over the valid subset. A lower pdbTM indicates greater structural distance from the training distribution. The pdbTM follows the definition of Anand et al. [2025] with two deviations: (i) we average over all valid samples across the full length grid, yielding a single value per model, whereas the original metric is reported separately for each length; (ii) structures are aligned on *C*3′ atoms (US-align default RNA mode) rather than *C*4′. Sample-level validity differences between a fixed DuetRNA checkpoint and each baseline are assessed using two-proportion z-tests with 95% confidence intervals; variability across training and inference seeds is reported separately as mean ± standard deviation.

### 5.2 Frame Representation Analysis

The representation analysis is independent of model training. Its static pool contains 11,497 processed RNAsolo structure units, while its multi-state pool contains 523 single-chain NMR entries and 31,432 relation groups. Interaction labels and geometric conventions follow established RNA base-pair and motif annotations [Leontis and Westhof, 2001, Sarver et al., 2008, Lu et al., 2015].

Base-anchored coordinates better preserve inter-residue organization. On the multi-state pool, Sugar-GS gives 13.35° rotational drift and 1.70 Å translational drift for canonical pairs. The deployed Base-Plane frame reduces these values to 7.78° and 0.61 Å. Base-Center reaches 7.76° and 0.47 Å, but Base-Plane is used because its origin is the glycosidic connection atom.

The reconstruction audit shows the complementary role of the sugar frame. Across 51,013 interaction pairs, Sugar-GS preserves backbone and next-residue bridge geometry but gives 6.21 Å base-channel RMSD. Combining Base-Plane with Sugar-GS lowers the base error to 1.58 Å while retaining the Sugar-GS backbone and bridge errors of 1.88 Å and 2.84 Å. The dual representation therefore combines the relation geometry of the deployed base channel with the reconstruction geometry of the sugar channel. Full results are given in Appendix B.

### 5.3 Protocol-Aligned Generation Results

#### Dual-frame structures are more designable

On the RNA-FrameFlow grid, DuetRNA improves IF validity from the value 41.00% of RNA-FrameFlow to 48.67%. This comparison uses the same RNAsolo corpus family, length grid, sample count, inverse-folding pipeline, and RhoFold evaluator. The 7.67 percentage-point gain is statistically significant. The 41.00% baseline is the published value; our own rerun of the official camera-ready checkpoint with the paper’s inference settings and default seed yields 35.5% IF validity and appears only in Table 3(b), where all models pass through the same evaluation pipeline for distribution-level comparison.

**Table 1:** Representative results from the frame analysis. Multi-state drift is evaluated in the relation-carrying channel of each configuration; for the dual-frame model this is the Base-Plane channel. Reconstruction uses base atoms from the base channel and sugar/backbone and bridge atoms from the Sugar-GS channel. Lower drift and reconstruction RMSD are better. The complete seven-frame sweep and category-level results are in Appendix B.

| Configuration | cWW multi-state drift |  | Reconstruction RMSD |  |  |
| --- | --- | --- | --- | --- | --- |
| | Rotation ($^{\circ}$) | Transl. ($\text{\AA}$) | Base ($\text{\AA}$) | Backbone ($\text{\AA}$) | Bridge ($\text{\AA}$) |
| Sugar-GS | 13.35 | 1.70 | 6.21 | **1.88** | **2.84** |
| Base-Plane | **7.78** | **0.61** | 5.85 | 2.98 | 4.06 |
| Base-Plane + Sugar-GS | **7.78** | **0.61** | **1.58** | **1.88** | **2.84** |

**Table 2:** Protocol-aligned comparisons. Each row matches the evaluation grid, sequence-source protocol, folding evaluator, and sample budget; the corpus family is RNAsolo. Method-specific preprocessing may still differ. DuetRNA values are for the headline checkpoint, the best of four independently trained runs (mean 45.68% ± 2.33%, Table 4). Confidence intervals refer to the difference in validity.

| Benchmark | Protocol | Evaluator | Baseline | DuetRNA | Difference | 95% CI | $p$ |
| --- | --- | --- | --- | --- | --- | --- | --- |
| RNA-FrameFlow grid | IF | RhoFold | 41.00% | **48.67%** | +7.67 pp | [2.04, 13.23] | 0.0076 |
| RiboFlow grid | IF | RhoFold | 34.70% | **44.33%** | +9.67 pp | [4.14, 15.11] | 0.0006 |
| RiboGen grid | GS | Boltz-1 | 34.17% | **38.50%** | +4.33 pp | [-1.11, 9.74] | 0.119 |

**Table 3:** Controlled mechanism and distribution analyses on RNAsolo. Panel (a) holds the split, training schedule, sampler, evaluator, and approximate parameter budget fixed. Panel (b) uses shared clustering and alignment pipelines. The RNA-FrameFlow rerun in Panel (b) uses its public checkpoint and default settings under IF pipeline. Lower reference-set pdbTM similarity indicates greater novelty.

| (a) Representation and coupling ablations |  |  |  |  |  |
| --- | --- | --- | --- | --- | --- |
| Model | IF validity |  | Difference from full |  |  |
| Full DuetRNA | 48.67% |  | – |  |  |
| Sugar-GS-only | 34.67% |  | -14.00 pp |  |  |
| Dual frame without relative-pose loss | 30.00% |  | -18.67 pp |  |  |
| (b) Validity, diversity, and novelty |  |  |  |  |  |
| Model | Validity | All-sample diversity | Valid-set diversity | Unique-valid yield | Reference-set pdbTM |
| RNA-FrameFlow rerun | 0.355 | 0.407 | 0.117 | 0.042 | 0.742 |
| Sugar-GS-only | 0.347 | 0.357 | 0.067 | 0.023 | 0.836 |
| Full DuetRNA | 0.487 | 0.268 | 0.134 | 0.065 | 0.822 |

#### The gain persists on a second inverse-folded benchmark

RiboFlow reports 34.70% under IF evaluation on its own length grid. Rerunning DuetRNA on the same grid and protocol gives 44.33%. This result supports the backbone-designability gain on a different length distribution. It does not compare the two models’ generated sequences.

#### Direct co-generation also achieves better performance

On the RiboGen GS benchmark with Boltz-1, DuetRNA reaches 38.50% and RiboGen reports 34.17%. The confidence interval includes zero, so we treat the difference as numerical rather than statistically established. DuetRNA also reaches 38.83% under GS with RhoFold on the RNA-FrameFlow grid.

### 5.4 Controlled Mechanism and Distributional Trade-offs

#### Both frames and their coupling contribute substantially

Removing the base-centered state reduces IF validity by 14.00 percentage points. Keeping both frames but removing relative-pose supervision reduces validity by 18.67 points. The first result shows that the base frame contributes beyond the sugar-centered representation used for backbone reconstruction. The second shows that two unconstrained frames are not sufficient. Their relative pose must also remain chemically coherent.

#### A higher yield of designable modes

DuetRNA has lower all-sample diversity *D*_all_, indicating a more concentrated overall generated distribution. After conditioning on valid samples, however, its valid-conditioned diversity *D*_valid_ is 0.134, compared with 0.117 for the RNA-FrameFlow rerun. Combined with its higher validity, this yields a higher unique-valid yield *Y*_UV_ (0.065 versus 0.042), indicating that DuetRNA produces more distinct designable structural modes per generation attempt. DuetRNA has higher similarity to the training set (0.822 vs. 0.742), however, indicating reduced novelty under this reference-set metric. The main benefit is therefore a higher yield of designable modes.

### 5.5 Robustness and Direct Geometry

#### The main result is not confined to one training run

Across the four training runs, mean IF validity is 45.68% ± 2.33%, and every run exceeds RNA-FrameFlow’s published 41.00%. GS performance has larger between-run variation. The sampler was selected once in the sampler sweep ablation and then fixed for all RNAsolo retrainings. Additional integration steps do not improve validity monotonically. The full sweep and the original training-budget analysis are reported in Appendix C.4.

#### Dual-frame generation improves most local geometry but introduces a specific linkage error

Relative to Sugar-GS-only, DuetRNA reduces steric clashes by 41.8% and improves the typical inter-residue phosphodiester geometry. DuetRNA’s North/South split (89.0%/11.0%) is close to the C3′-endo/C2′-endo proportions observed in experimental RNA (roughly 80–90% North), whereas the Sugar-GS-only ablation over-collapses onto C3′-endo (93.9%/6.1%). Its *O*3′-*P* RMSD remains slightly higher because a small catastrophic tail dominates the squared error despite a better outlier rate and P99 deviation. The main local weakness occurs at the glycosidic *C*1′-*N* 9*/N* 1 bond. DuetRNA reconstructs the sugar and base atoms from two predicted frames, while Sugar-GS-only places both through one fixed rigid template. The dual-frame model therefore improves packing and chain geometry but loses exact glycosidic linkage.

### 5.6 Qualitative Analysis

We complement the quantitative metrics with a visual inspection of representative generated structures and their geometric quality. Figure 3 shows eight samples spanning lengths 40-150 nt, each displaying the generated structure (rainbow ribbon) overlaid with the RhoFold prediction of its designed sequence (dark grey) and labeled with the sample length and the corresponding scTM, scRMSD, and scGDT values. An additional panel illustrates the dominant geometric failure modes: chain breaks, local base-base clashes, and an excessively long loop (Figure 4).

**Figure 3:**
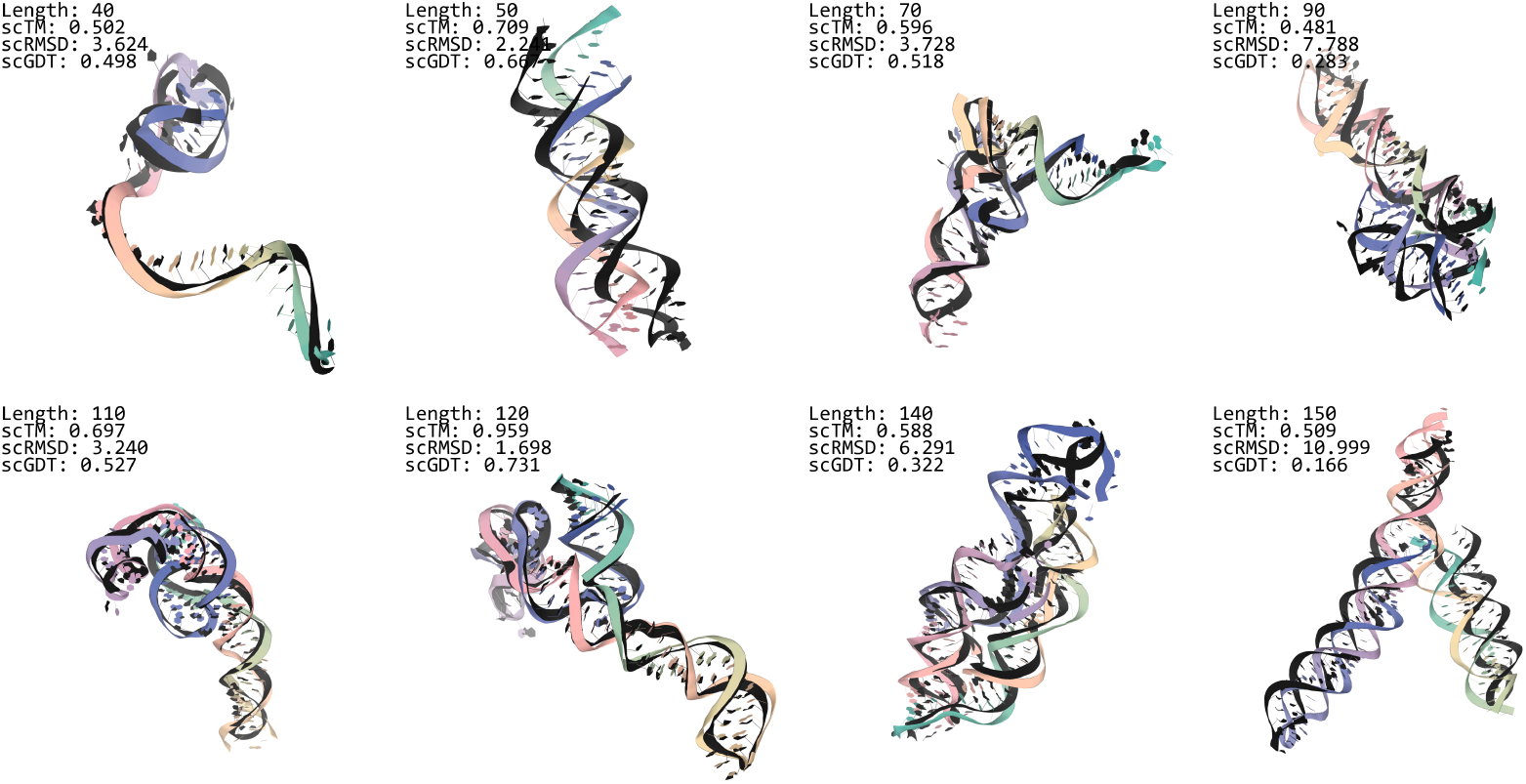
Qualitative analysis. For each representative sample (40-150 nt), the generated structure (rainbow ribbon) is overlaid with the RhoFold prediction of its designed sequence (dark grey); each panel is labeled with the sample length and its scTM, scRMSD, and scGDT values.

**Figure 4:**
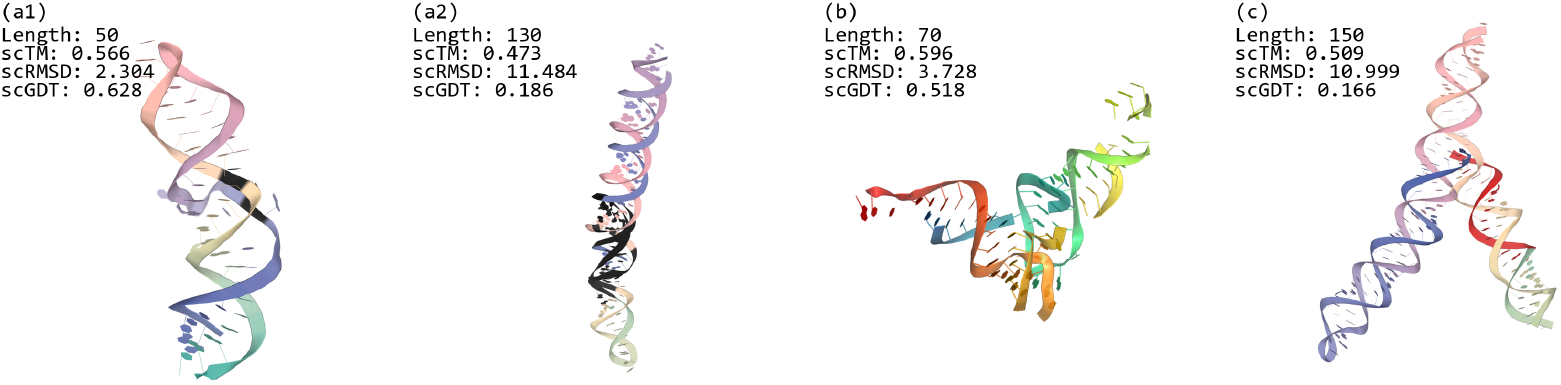
Geometric failure modes in generated samples. (*a*_1_) Base-base clash in a 50-nt sample: the clashing base pair and the adjacent backbone contacts (residues 12, 13, and 36) are colored black. (*a*_2_) A more severe clash in a 130-nt sample (21.3 clashes per 100 atoms): a contiguous stretch of clashing residues in the middle of the chain is colored black. (b) Chain breaks in a 70-nt sample: the generated structure is split into seven fragments, five of them shorter than ten residues. (c) Excessive loop in a 150-nt sample: a 16-nucleotide unpaired segment (red) forms an unusually extended loop region.

In Figure 3, for the 120-nt sample, the generated structure closely tracks the RhoFold prediction (scTM 0.959, scRMSD 1.70 Å), forming a single continuous chain with no steric clashes and no chain breaks. The 40-nt and 110-nt samples preserve the overall topology, with deviations concentrated in loop regions, in line with their intermediate scTM values. These observations are consistent with the quantitative observation that geometric quality is best in the 120-nt regime and remains intact for moderate lengths.

Figure 4 shows three frequent failure modes. First, severe base-base clashes are confined to a minority of samples but concentrate at short lengths, where the two ends of a collapsed chain interpenetrate (the most severe sample exceeds 115 clashes per 100 atoms); the 50-nt and 130-nt examples contain local clusters of clashing residues, colored black (Figure 4a). Second, fragmentation is the dominant geometric failure: across the 600-sample batch, chain breaks affect up to 94% of 70-nt samples, and the highlighted 70-nt sample (Figure 4b) is split into seven fragments, five of them shorter than ten residues. Third, the 150-nt sample contains a 16-nucleotide unpaired run (Figure 4c). Natural hairpin-loop lengths are strongly bimodal, with tetraloops (4 nt) most frequent and heptaloops (7 nt) second, and loops longer than 7 nt occurring much less often [Danaee et al., 2018]; a contiguous 16-nt unpaired segment inside a compact 150-nt fold is therefore atypical and suggests distorted secondary structure, although unpaired runs of this length do occur in natural RNAs such as pre-miRNA apical loops and multi-helix junctions.

Notably, scTM and scRMSD do not strongly penalize disconnectedness: the 70 nt sample in Figure 4b attains a scTM of 0.596 and a scRMSD of 3.73 despite the high break rate. We also note that the reference is a RhoFold prediction rather than an experimental structure, so the visual agreement measures consistency with an independent folding model rather than ground-truth correctness.

Taken together, the visual evidence corroborates the quantitative results: DuetRNA produces geometrically sound, self-consistent designs in its favorable length regime, while the failure modes concentrate at lengths away from this regime. The RNAsolo training set is unbalanced across sequence lengths, and for samples whose lengths are underrepresented, the learned physical constraints are insufficient, which manifests as chain breaks, local clashes, and distorted secondary structure.

## 6 Conclusion

In this work, we introduce DuetRNA, a dual-frame SE(3) flow-matching framework for RNA sequence-structure co-generation. Motivated by a systematic analysis of seven single-frame candidates across 11,497 static chains and 31,432 multi-state relation groups, DuetRNA represents each nucleotide with coupled base- and sugar-centered frames, enabling residue-level organization and intra-residue backbone reconstruction geometry to be generated within a unified flow.

Our results show that DuetRNA with this factorization yields strong folding-based self-consistency, achieving 48.67% IF scTM-validity on the RNA-FrameFlow grid and 38.83% GS scTM-validity with RhoFold. Removing the base frame or the relative-pose objective substantially reduces IF validity, and direct atom23 analyses show improved steric packing, sugar-ring closure, and typical phosphodiester geometry relative to a capacity-matched Sugar-GS-only model. These results support a geometric view of RNA generation in which base-centered frames capture nucleobase-mediated pairing, stacking, and long-range organization, while sugar-centered frames preserve the degrees of freedom needed for atomistic backbone reconstruction.

The present evidence is limited to unconditional generation of 40-150 nt RNAs under computational self-consistency and coordinate-level analyses. It does not establish biochemical function, experimental viability, or generalization to longer RNAs. Valid samples are also closer to the training corpus, and two-frame reconstruction weakens glycosidic-linkage accuracy. Absolute stereochemical quality remains imperfect, with substantial clash and linkage-outlier rates and length-dependent chain breaks. Future work should address these limitations before extending the framework to broader tasks like function-conditioned design and multi-conformation generation.

## Code and Data Availability

The official DuetRNA repository is https://github.com/XjunLi/DuetRNA. The raw RNA structures are drawn from the publicly available RNAsolo and RNA3DB resources [Adamczyk et al., 2022, Szikszai et al., 2024]; no new experimental data were collected. External predictors and analysis tools remain subject to their original licenses and distribution terms.

## A Method Details

### A.1 Model Architecture

#### A.1.1 Dual-Frame IPA-Transformer Trunk

Our geometric trunk extends the Invariant Point Attention (IPA) framework introduced by AlphaFold2 [Jumper et al., 2021] to a dual-frame setting. IPA’s core idea is to process 3D residue frames through SE(3)-invariant attention by projecting local coordinate differences into invariant spatial features, enabling the network to reason about relative geometry without overfitting to global pose.

##### Shared pair representation

Each geometric stream is initialized from shared timestep and positional embeddings together with a frame-type embedding distinguishing base and sugar objects. The trunk constructs three pair channels: a base-view pair representation from base-node features and base translations, a sugar-view pair representation from sugar-node features and sugar translations, and a bridge-view pair representation from a per-residue summary of the base and sugar states together with their translation difference. These channels are fused into a shared pair tensor used by both geometric streams.

##### Dual geometric streams

At block *k*, the trunk maintains 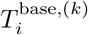 and 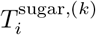 together with their corresponding feature streams. Two invariant point attention modules process the current base and sugar poses in parallel, followed by per-stream transitions and rigid updates

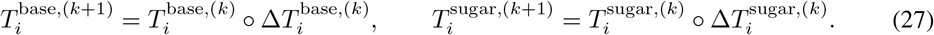

##### Object-token mixing and dynamic edge refinement

To enable explicit cross-object communication, the residue-wise base and sugar tokens are periodically concatenated into a sequence of 2*L* object tokens and processed by a joint transformer encoder. Between blocks, residue-level summaries from the current base and sugar streams are written back into the shared pair tensor through an edge transition, so the pair representation evolves jointly with trunk depth.

##### Self-conditioning

The pair stack also receives distograms computed from self-conditioned translations. During training, with probability 0.5, a stop-gradient forward pass provides predicted base and sugar translations that are fed back into the pair embedders. During sampling, the previous-step predictions play the same role.

##### Sequence and Torsion Heads

Let 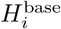 and 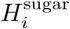 denote the final residue features of the two streams. The sequence head reads the final base token together with pooled pair context from the base-view, sugar-view, and fused pair representations. The torsion head reads the final sugar feature together with the initial sugar embedding.

### A.2 Auxiliary Loss Definitions

#### Soft Base-Template Loss

For each candidate nucleotide *c* ∈ *C*, let 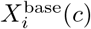 be the base-template atomic coordinates of nucleotide *c* placed in the predicted base frame at residue *i*. Given predicted categorical probabilities *π*_*i*_(*c*), the probability-weighted base-template coordinates are

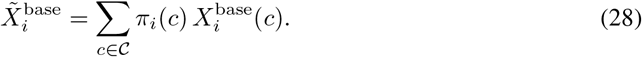

The corresponding auxiliary loss is

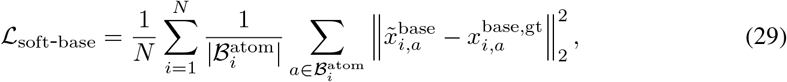

where 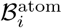 is the atom23 base-atom subset for residue *i*. This term gives the sequence probabilities structural feedback through base-template coordinates; it does not replace the categorical cross-entropy objective.

#### Chain and Clash Auxiliary Losses

Let ℰ denote the valid adjacent-residue set for which both 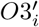 and *P*_*i*+1_ are present. The chain-continuity auxiliary supervises the decoded adjacent 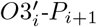 bridge:

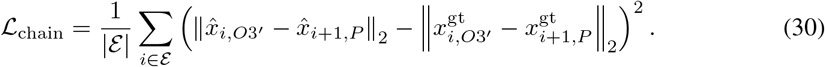

Let *V* denote the set of valid residues, *N*_valid_ = |*V*|, and let *A*_*i*_ denote the valid atom23 heavy atoms of residue *i*. Define the set of nonlocal inter-residue atom pairs as

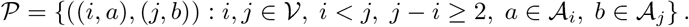

The clash auxiliary loss is

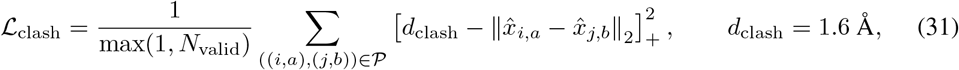

where [*z*]_+_ = max(0, *z*). We use 1.6 Å as a conservative severe-clash threshold, approximately matching the canonical inter-residue O3’-P phosphodiester bond length (1.602 Å). After excluding the actual covalent O3’-P linkage, any inter-residue heavy-atom pair closer than this scale is treated as physically implausible self-intersection. This is deliberately looser than a VdW-based steric cutoff, so it penalizes only severe overlaps rather than legitimate tight RNA packing, hydrogen-bonding, or stacking contacts.

### A.3 Inference

Inference keeps two rigid objects for every residue throughout the whole trajectory. We first sample independent source chains 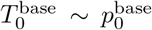 and 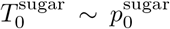. At step *t*, one coupled model evaluation takes 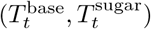 as input and predicts terminal frames 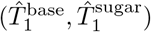. Both channels are then updated in parallel:

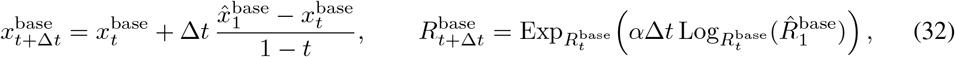

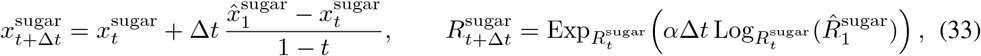

where *α* is the rate parameter of the exponential rotation sampling schedule.

If self-conditioning is enabled, the previous-step predicted base and sugar translations are fed back into the pair representation. After the final integration step, the model produces sequence probabilities and intra-residue torsion estimates from the terminal base and sugar features. Then we choose the sequence by

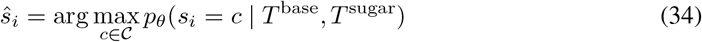

and complete atom23 coordinates from the terminal pair of frames:

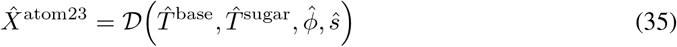

to obtain the final atom23 structure.

## B Frame Representation Analysis: Extended Results

This appendix provides the extended diagnostics underlying the frame representation analysis of §5.2. The main text reports the two core findings: base anchoring better captures inter-residue relation geometry, while a separate sugar/backbone frame is needed for local reconstruction. Here we report the candidate-frame catalog and deployed coordinate definitions, the canonicalization and continuity screen (§B.2), the static relation-separation analysis (§B.3), the per-relation multi-state breakdown including noncanonical pairs and stacking (§B.4), and the single-frame and dual-frame reconstruction sweep (§B.5).

### B.1 Candidate Frame Catalog

We audit seven residue-level single-frame candidates spanning sugar/backbone and base-anchored constructions. The display names below are used consistently in the paper; the corresponding implementation identifiers are shown in parentheses.

#### Base family

- **Base-Plane** (`base_plane_anchor`): a chemically anchored base-plane frame. The origin is the glycosidic connection atom (*N* 9 for purines and *N* 1 for pyrimidines). The orientation is defined by the fitted nucleobase plane normal and an in-plane chemical axis (*C*4 → *C*8 for purines, *C*4 → *C*2 for pyrimidines).
- **Base-Inertial** (`base_inertial`): an inertial frame over all nucleobase heavy atoms, with residue-type-specific sign and quadrant disambiguation. The origin remains the glycosidic connection atom.
- **Base-Center** (`base_center_standard`): the base-origin control. It uses the same inertial orientation family as Base-Inertial, but moves the origin to the mass centroid of the nucleobase heavy atoms.

#### Sugar/backbone family

- **Sugar-GS** (`baseline_geom`): the legacy three-atom Gram-Schmidt frame built from (*O*4′, *C*4′, *C*3′) with origin at *C*4′, following the axis convention used in RNA-FrameFlow [Anand et al., 2025].
- **Sugar-Inertial-4** (`sugar4_inertial`): a four-atom mass-weighted inertial frame over (*O*4′, *C*4′, *C*3′, *C*5′) with origin at *C*4′ and anchor-based sign disambiguation.
- **Sugar-Inertial-6** (`sugar6_inertial`): a six-atom inertial variant over (*C*1′, *C*2′, *C*3′, *C*4′, *O*4′, *C*5′) with origin at *C*4′.
- **Sugar-Ring** (`sugar5_ring_axis`): a five-membered sugar-ring frame over (*O*4′, *C*1′, *C*2′, *C*3′, *C*4′), with ring-plane orientation and a sugar-ring centroid origin.

#### B.1.1 Deployed Frame Definitions

The deployed dual-frame model uses Sugar-GS for 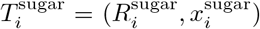 and Base-Plane for 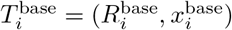. We give their exact coordinate definitions here.

##### Sugar-GS

Let *x*_*O*4_*′, x*_*C*4_*′, x*_*C*3_′ ∈ ℝ ^3^ be the corresponding atom coordinates. We set the origin at *C*4′,

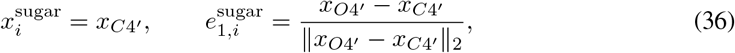

orthogonalize the *C*3′ direction against 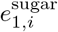,

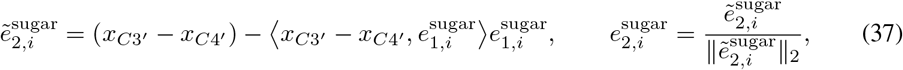

and complete a right-handed frame,

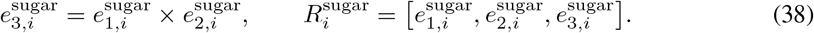

##### Base-Plane

Let ℬ_*i*_ denote the heavy-atom set of the nucleobase, excluding sugar atoms, and let 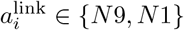 be the glycosidic connection atom. The origin is

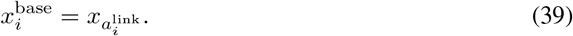

We fit a plane to {*x*_*a*_ : *a* ∈ ℬ_*i*_} and denote its unit normal by *ñ*_*i*_. To resolve the normal sign, we use the residue-type-specific reference normal

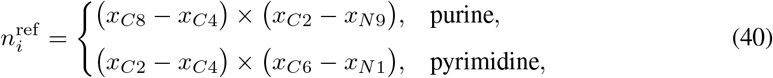

and set

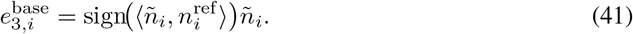

The in-plane chemical axis is

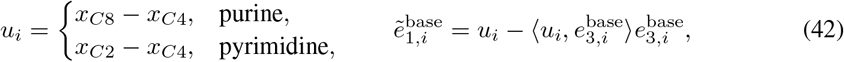

which gives

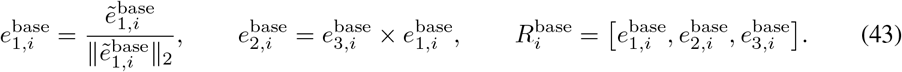

### B.2 Frame Canonicalization and Continuity Screening

Before comparing downstream relation or reconstruction metrics, each frame must provide a usable SE(3) supervision target. We therefore first apply a canonicalization and continuity gate. A candidate passes only if its valid rate is at least 0.95, the maximum deterministic rebuild error is at most 10^−5^ degrees, and the fraction of perturbation cases with a rotation jump above 150° is zero. The perturbation suite includes micro sugar-pucker, *χ*-rotation, phosphate-swing, and bridge-motion probes.

Sugar-Inertial-4 is therefore not used as a formal relation-stability frame in the main evidence chain. Its average continuity is not poor, but the small nonzero catastrophic-jump tail makes it unsafe as a smooth flow-matching label. We retain it only as an exploratory reconstruction diagnostic when explicitly marked as such.

### B.3 Static Relation Separation

On the full static pool of 11,497 processed RNAsolo structure units, 1,765 nonredundant representatives, and 1,045 motif-rich chains, we examine which frame family more readily separates relation types and which more tightly clusters identical relation instances. We run 20 independent sub-experiments, structured as one full set and three filtered subsets (representative, motif-rich, stem-like), across five relation categories (canonical cWW pairs, noncanonical pairs, stacking, base-phosphate, and base-backbone). Canonical cWW pairs include Watson-Crick AU/UA and GC/CG pairs plus GU/UG wobble pairs. In 11 of the 20 subset-relation combinations, the frame with the best inter-group/intra-group distance ratio belongs to the sugar/backbone family while the frame with the best 1-NN retrieval accuracy belongs to the base family. This result implies that frames rooted in sugar/backbone atoms spread different relation categories farther apart, whereas frames rooted in base atoms group geometrically identical instances closer together. No single frame family achieves the best result in every channel. Table 6 reports the family-level split pattern.

**Table 4:** Training robustness and direct atom23 geometry. The headline checkpoint is evaluated with the fixed main sampler; values after ± are standard deviations over three inference seeds for the corresponding independently trained checkpoint. Geometry is evaluated on the 600 generated structures of the main length grid (40–150 nt, 50 per length); the length-stratified breakdown is given in Appendix C.7. Two unbonded heavy atoms *i, j* clash if *r*_*ij*_ *< v*_*i*_ + *v*_*j*_ − 0.6 Å, normalized per 100 heavy atoms, following Anand et al. [2025].

| (a) Independent training runs |  |  |
| --- | --- | --- |
| Checkpoint | IF validity | GS validity |
| Headline checkpoint | 48.67% | 38.83% |
| Seed 123 | 43.11% ± 0.77% | 36.33% ± 1.33% |
| Seed 456 | 44.89% ± 1.55% | 39.94% ± 1.35% |
| Seed 789 | 46.06% ± 2.10% | 31.61% ± 2.91% |
| (b) Direct coordinate analysis |  |  |
| Metric | Full DuetRNA | Sugar-GS-only |
| Clashes per 100 atoms | **26.55** | 45.58 |
| North / South pucker | 89.0% / 11.0% | 93.9% / 6.1% |
| Mean pucker amplitude | 34.92° | 38.62° |
| Sugar-ring closure RMSD | **0.099 Å** | 0.176 Å |
| $C1'-N9$ RMSD | 0.417 Å | **0.003 Å** |
| $C1'-N1$ RMSD | 0.484 Å | **0.003 Å** |
| Inter-residue $O3'-P$ outlier rate | **38.6%** | 51.4% |
| Inter-residue $O3'-P$ P99 deviation | **0.857 Å** | 4.990 Å |
| Inter-residue $O3'-P$ RMSD | 1.267 Å | **1.045 Å** |

**Table 5:** Frame canonicalization and continuity screening. The gate is strict on catastrophic rotation jumps: any nonzero *>* 150° jump rate fails the candidate.

| Frame | Valid rate | Median cont.<br>($^\circ$) | P95 cont.<br>($^\circ$) | Jump<sub>150</sub> | Gate |
| --- | --- | --- | --- | --- | --- |
| Sugar-GS (baseline_geom) | 1.00000 | 1.88 | 5.00 | 0 | pass |
| Sugar-Inertial-4 (sugar4_inertial) | 0.99982 | 1.16 | 22.36 | $9.8 \times 10^{-4}$ | fail |
| Sugar-Inertial-6 (sugar6_inertial) | 0.99969 | 0.59 | 5.00 | 0 | pass |
| Sugar-Ring (sugar5_ring_axis) | 0.99987 | 0.89 | 5.00 | 0 | pass |
| Base-Plane (base_plane_anchor) | 0.99584 | 0.00 | $1.7 \times 10^{-6}$ | 0 | pass |
| Base-Inertial (base_inertial) | 0.99584 | 0.00 | $1.7 \times 10^{-6}$ | 0 | pass |
| Base-Center (base_center_standard) | 0.99584 | 0.00 | $1.7 \times 10^{-6}$ | 0 | pass |

**Table 6:**
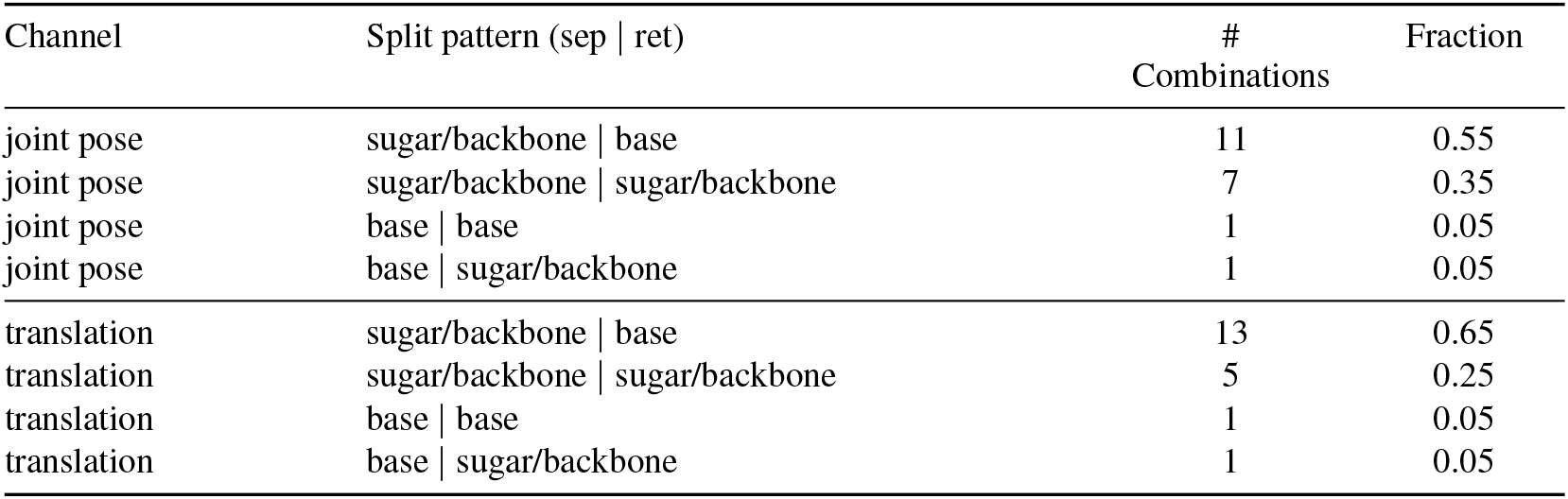
Family-level split pattern on the static pool. Each row reports the number of subset-relation combinations (out of 20) in which the indicated pattern holds: “sep” is the family that better separates relation types, and “ret” is the family that more tightly retrieves identical relation instances.

For a residue pair (*i, j*), the inter-residue relation is represented by

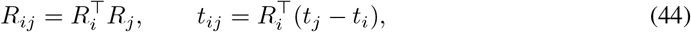

where *R*_*i*_ and *t*_*i*_ are the rotation and translation of residue *i*.

These counts are descriptive evidence of two distinct representational roles in RNA residue geometry, not formal statistical claims.

### B.4 Multi-State Stability: Per-Relation Breakdown

Due to the flexibility of RNA, the same molecule may adopt different conformations. A frame that separates relation types in a static snapshot does not necessarily keep the same relation stable across conformational states. NMR and other multi-model entries therefore provide a natural test bed for measuring drift across plausible states of the same molecule.

Here we extend the multi-state analysis of §5.2 to the five paper-facing frames used in the completed external run: Sugar-GS, Sugar-Inertial-6, Sugar-Ring, Base-Plane, and Base-Center. The pool consists of 523 single-chain multi-model entries (496 tier-A NMR plus 27 other multi-model structures), comprising 31,432 relation groups, 984,382 normalized relation rows, and 336 high-coverage relation keys. Variation reflects a combination of conformational flexibility and experimental uncertainty.

For each key, *orientation drift* Δ*R* (median geodesic angle from the medoid rotation) and *translation drift* Δ*T* (median Euclidean distance from the medoid translation) are computed under each candidate frame. A key is *base-like* in a channel if the best base-family frame outperforms all sugar/backbone-family frames.

Canonical cWW pairs strongly favor base-anchored frames in both rotation and translation channels. Noncanonical pairs favor base-anchored frames at the aggregate level, but remain key-heterogeneous: among 191 high-coverage noncanonical relation keys, 76 keys exhibit a sugar/backbone-like rotation pattern with a base-like translation pattern, while by evaluation weight 74% of pairs are base-like in both channels. Stacking is the most heterogeneous: by evaluation weight, 93.6% of stacking pairs favor base-like rotation stability, while 44.6% of stacking translations remain sugar/backbone-like; by key count, 72.7% of stacking keys are base-like in rotation. The multi-state evidence therefore identifies canonical cWW pairing as cleanly base-governed and stacking as a relation in which orientation and translation behave differently.

Table 9 reports, for each category, the fraction of keys whose lowest drift comes from a base-family frame. For canonical cWW pairing, a base-family frame achieves the lowest drift for every key in both channels, consistent with the physical organization of Watson-Crick and wobble pairing at the nucleobase level. Noncanonical pairing and stacking are more heterogeneous: only 38% of noncanonical keys have their lowest orientation drift from a base-family frame, yet 68% have their lowest translation drift from a base-family frame; stacking shows 73% of keys favoring a base-family frame in orientation and 68% in translation. Thus, no single frame family yields the lowest drift in both channels for every relation type.

**Table 7:**
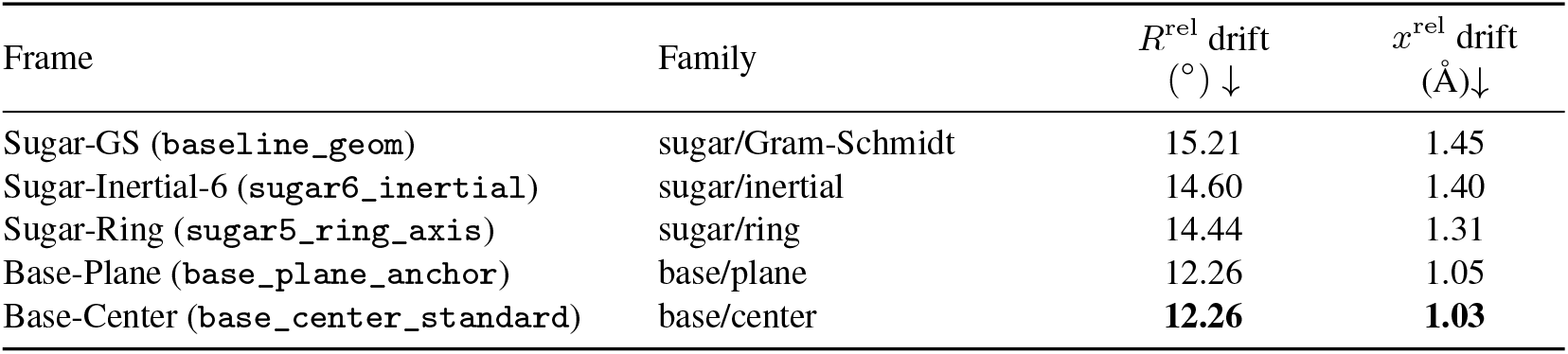
Aggregate multi-state stability over all 31,432 relation groups. Lower is better.

| Frame | Family | $R^{\text{rel}}$ drift<br>($^{\circ}$) ↓ | $x^{\text{rel}}$ drift<br>($\text{\AA}$) ↓ |
| --- | --- | --- | --- |
| Sugar-GS (baseline_geom) | sugar/Gram-Schmidt | 15.21 | 1.45 |
| Sugar-Inertial-6 (sugar6_inertial) | sugar/inertial | 14.60 | 1.40 |
| Sugar-Ring (sugar5_ring_axis) | sugar/ring | 14.44 | 1.31 |
| Base-Plane (base_plane_anchor) | base/plane | 12.26 | 1.05 |
| Base-Center (base_center_standard) | base/center | **12.26** | **1.03** |

**Table 8:**
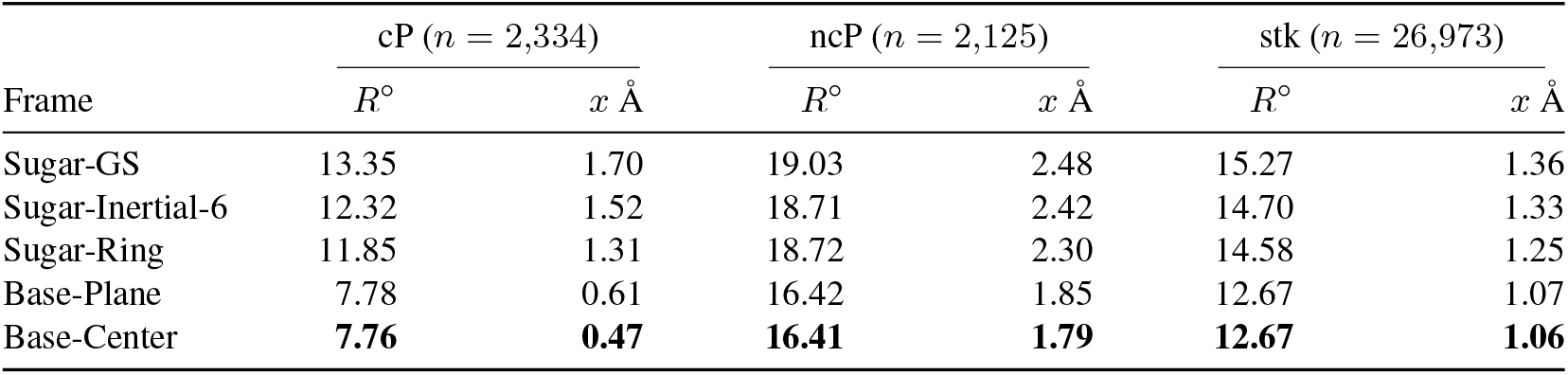
Per-relation multi-state stability across the five external-run frames. Lower is better. “cP” = canonical cWW pairs (*n* = 2,334), “ncP” = noncanonical pairs (*n* = 2,125), and “stk” = stacking (*n* = 26,973).

**Table 9:** Fraction of high-coverage relation keys whose lowest drift is achieved by a base-family frame.

| Relation category | Keys | Orientation | Translation |
| --- | --- | --- | --- |
| Canonical cWW pairs | 6 | 100% | 100% |
| Noncanonical pairs | 191 | 38% | 68% |
| Stacking | 139 | 73% | 68% |

### B.5 Reconstruction Path: Full Single-Frame and Dual-Frame Sweep

The main text reports the core reconstruction tradeoff. Here we report the full reconstruction-path sweep on 51,013 interaction pairs for the gate-passing single frames supported by this path and the three dual configurations that pair a base-anchored frame with the Gram-Schmidt Sugar-GS frame. Sugar-Inertial-4 is excluded from this formal table because it failed the screening gate in Table 5.

**Table 10:** Reconstruction-path comparison on 51,013 interaction pairs. RMSD_base_ measures atoms reconstructed in the base channel; RMSD_bb_ and RMSD_bridge_ measure atoms reconstructed in the sugar/backbone channel and the next-residue *O*3′ → *P* bridge.

| Configuration | $\text{RMSD}_{\text{base}}$<br>(Å)↓ | $\text{RMSD}_{\text{bb}}$<br>(Å)↓ | $\text{RMSD}_{\text{bridge}}$<br>(Å)↓ | Clash rate<br>↓ |
| --- | --- | --- | --- | --- |
| *Single-frame configurations* |  |  |  |  |
| Sugar-GS | 6.21 | 1.88 | 2.84 | $4.9 \times 10^{-4}$ |
| Sugar-Ring | 5.37 | 1.85 | 2.75 | $5.5 \times 10^{-2}$ |
| Sugar-Inertial-6 | 7.12 | 1.96 | 2.30 | $8.3 \times 10^{-4}$ |
| Base-Plane | 5.85 | 2.98 | 4.06 | $4.9 \times 10^{-4}$ |
| Base-Inertial | 5.60 | 2.34 | 2.91 | $2.1 \times 10^{-3}$ |
| Base-Center | 5.61 | 3.20 | 3.74 | $2.4 \times 10^{-3}$ |
| *Dual-frame configurations with Sugar-GS local frame* |  |  |  |  |
| Base-Plane + Sugar-GS | 1.58 | 1.88 | 2.84 | $4.8 \times 10^{-4}$ |
| Base-Inertial + Sugar-GS | 1.52 | 1.88 | 2.84 | $4.8 \times 10^{-4}$ |
| Base-Center + Sugar-GS | 1.09 | 1.88 | 2.84 | $4.9 \times 10^{-4}$ |

Among single-frame choices, base-anchored frames preserve the relation geometry identified in the preceding sections but suffer from poor local sugar/backbone reconstruction; sugar/backbone frames give better local reconstruction but cannot accurately recover base atoms. The dual configurations decouple these roles: replacing the global frame with a base-anchored channel substantially reduces base-channel RMSD while retaining the Sugar-GS local reconstruction path.

Overall, the three analyses converge on the same conclusion: selecting any one single frame as a universal representative is inadequate for RNA. Base-anchored frames best capture inter-residue relation geometry (§B.3, §B.4) but do not by themselves solve local sugar/backbone reconstruction (§B.5); sugar/backbone-anchored frames show the complementary behavior. The dual-frame design resolves this by assigning each role to a dedicated channel: a global base frame for relation-bearing geometry and a sugar frame for reconstruction-bearing geometry (§4.1).

## C Experiment Details and Additional Analyses

### C.1 Benchmark Provenance

**Table 11:**
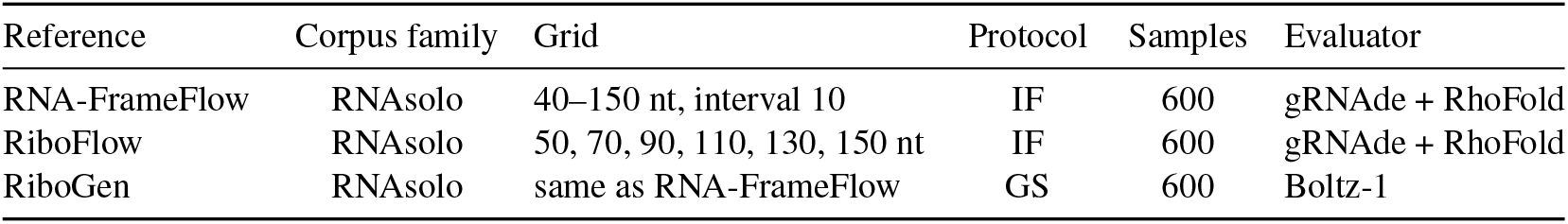
Provenance of the primary generation comparisons. DuetRNA is rerun for every row. Published baseline values retain method-specific preprocessing choices that cannot be made identical without the original training artifacts. Throughout this work, “RhoFold” denotes the RhoFold model and checkpoint used in the RNA-FrameFlow evaluation pipeline [Shen et al., 2022, Anand et al., 2025], rather than RhoFold+ [Shen et al., 2024].

IF and GS require different model outputs. IF can evaluate a backbone-only generative model because an external inverse-folding model supplies sequences. GS requires a model-generated sequence and therefore evaluates the generated pair directly. RhoFold and Boltz-1 also define different evaluators. We consequently avoid best-value highlighting across protocols or folding models.

### C.2 Auxiliary Objective Ablation

This development ablation uses the RNA3DB training split, a 120K training budget, and the fixed sampler (*N*_*T*_ = 50, *α* = 10). This ablation precedes the sampler sweep, so the fixed sampler follows RNA-FrameFlow’s settings. All variants keep the core frame, sequence, torsion, and atom23 objectives. The auxiliary losses are defined in Appendix A.2.

**Table 12:**
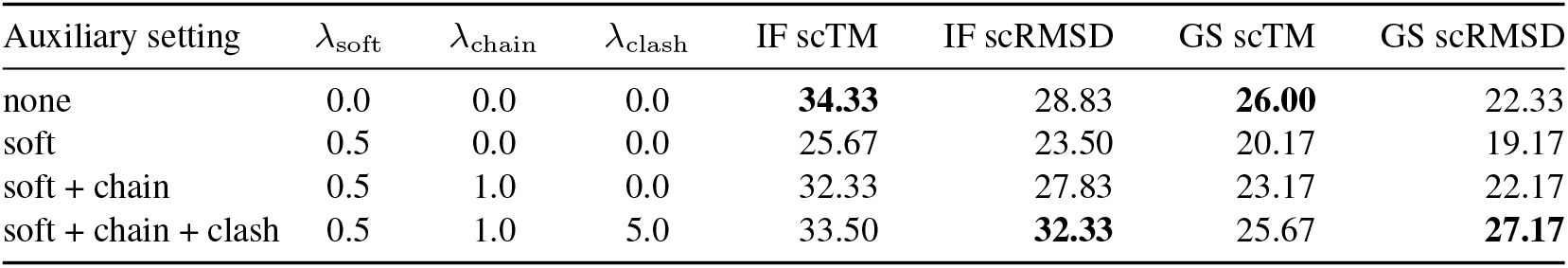
Auxiliary objective ablation on the RNA3DB development setting. All rows use 600 generated structures. Evaluated by IF scTM, IF scRMSD, GS scTM and GS scRMSD validity.

| Auxiliary setting | $\lambda_{\text{soft}}$ | $\lambda_{\text{chain}}$ | $\lambda_{\text{clash}}$ | IF scTM | IF scRMSD | GS scTM | GS scRMSD |
| --- | --- | --- | --- | --- | --- | --- | --- |
| none | 0.0 | 0.0 | 0.0 | **34.33** | 28.83 | **26.00** | 22.33 |
| soft | 0.5 | 0.0 | 0.0 | 25.67 | 23.50 | 20.17 | 19.17 |
| soft + chain | 0.5 | 1.0 | 0.0 | 32.33 | 27.83 | 23.17 | 22.17 |
| soft + chain + clash | 0.5 | 1.0 | 5.0 | 33.50 | **32.33** | 25.67 | **27.17** |

The core objective gives the highest scTM validity. The soft template loss alone reduces both IF and GS performance. Chain and clash terms recover part of this loss, and the clash term improves scRMSD validity. These objectives therefore change the operating point rather than improving all criteria together.

### C.3 Hyperparameter Ranges

**Table 13:** Core objective weights and auxiliary-loss ranges.

| Symbol | Reported value(s) | Ablation range | Role |
| --- | --- | --- | --- |
| $\tau$ | 0.25 | [0.1, 0.5] | time gate for decoded-geometry supervision |
| $\lambda_{\text{Seq}}$ | 1.0 | [0.5, 2.0] | nucleotide cross-entropy |
| $\lambda_{\text{Loc}}$ | 1.0 | [0.25, 2.0] | torsion and atom23 reconstruction |
| $\lambda_{\text{soft}}$ | 0 or 0.5 | [0, 1.0] | soft base-template auxiliary |
| $\lambda_{\text{chain}}$ | 0 or 1.0 | [0, 2.0] | adjacent $O3'-P$ continuity |
| $\lambda_{\text{clash}}$ | 0 or 5.0 | [0, 5.0] | steric clash auxiliary |

### C.4 Sampler and Training-Budget Sensitivity

The sampler was selected on the original RNA3DB development run and then frozen for all RNAsolo retrainings. *N*_*T*_ is the number of ODE integration steps, and *α* is the rate parameter of the exponential rotation sampling schedule.

**Table 14:** Inference schedule on the 120K RNA3DB development model. All rows use the same weights, RhoFold evaluator, and 600 generated structures.

| $N_T$ | $\alpha$ | IF | | GS | |
| --- | --- | --- | --- | --- | --- |
|  |  | scTM validity | Diversity | scTM validity | Diversity |
| 50 | 10 | 32.00 | **0.585** | 24.67 | **0.587** |
| 100 | 20 | **40.67** | 0.443 | **31.17** | 0.442 |
| 200 | 10 | 37.83 | 0.487 | 27.00 | 0.485 |
| 200 | 20 | 38.00 | 0.477 | 28.50 | 0.485 |
| 300 | 10 | 36.00 | 0.485 | 27.50 | 0.485 |
| 300 | 20 | 38.17 | 0.433 | 28.83 | 0.438 |

The highest validity occurs at (*N*_*T*_ = 100, *α* = 20), while the shortest trajectory gives the highest diversity. Increasing the number of integration steps does not improve validity monotonically. The selected setting is therefore an empirical validity-oriented operating point rather than a general benefit from additional computation.

**Table 15:**
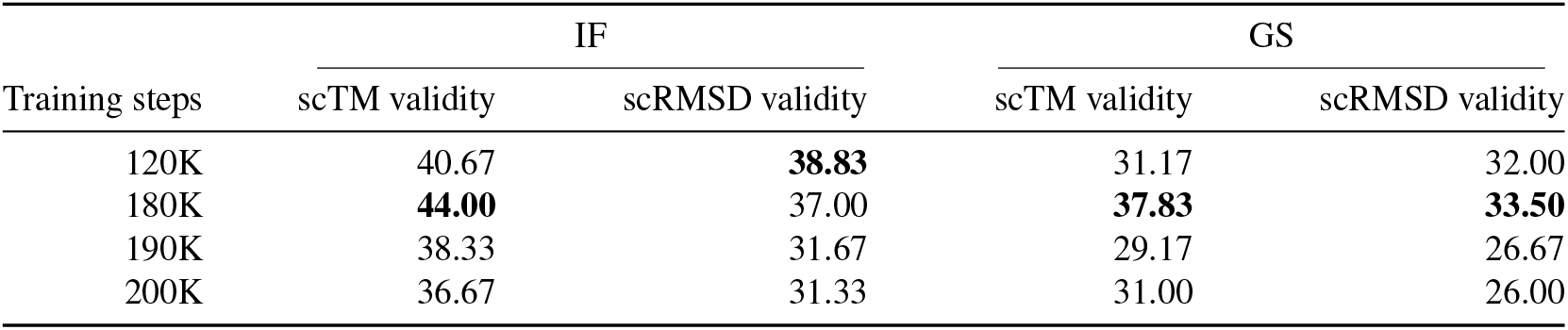
Training-budget sensitivity on the RNA3DB development run with the fixed (*N*_*T*_ = 100, *α* = 20) sampler.

Performance peaks at an intermediate checkpoint and then decreases. This observation motivated the independent RNAsolo training runs in Table 4. Their results show that the matched IF gain persists across optimization seeds even though checkpoint and GS sensitivity remain.

### C.5 Statistical Details for Matched Comparisons

All primary validity metrics are binomial proportions over 600 generated structures. The RNA-FrameFlow row compares 292*/*600 DuetRNA samples with 246*/*600 baseline samples. The Ri-boFlow row compares 266*/*600 with the 208*/*600 count implied by the rounded published rate. The RiboGen row compares 231*/*600 with the 205*/*600 count implied by 34.17%. Two-proportion tests give *p* = 0.0076, *p* ≈ 0.0006, and *p* = 0.119, respectively. The RiboFlow interval and test should be read as approximate because the exact unrounded baseline count was not reported. Its published 34.70% is rounded and implies 208 valid samples out of 600.

Intervals are Newcombe’s hybrid score interval for the difference of two independent proportions; *p*-values are two-sided pooled two-proportion *z*-tests.

### C.6 Scope of Direct Geometry Evaluation

The atom23 audit compares DuetRNA with Sugar-GS-only because both models produce the same complete atom set under matched capacity, data, training, and decoding pipelines. RNA-FrameFlow produces a reduced backbone representation and cannot support the same base-atom and glycosidic-linkage audit. The reported checks cover clashes, sugar pucker, glycosidic bonds, and the inter-residue *O*3′–*P* bond. They do not constitute a formal MolProbity score. Hydrogen-complete refinement, comprehensive bond-angle analysis, glycosidic *χ* analysis, biochemical assays, and experimental structure determination remain outside the current evaluation.

### C.7 Length-Stratified Analysis

We complement the aggregate geometry numbers of Table 4 with a per-length breakdown on the main generation grid: the 600 structures spanning 40-150 nt at intervals of 10 (50 per length), evaluated under the IF protocol (gRNAde + RhoFold) of §5.1. Fig. 5 reports the per-length scTM distribution together with the per-length validity rate under the single criterion scTM ≥ 0.45, and Fig. 6 the per-length scGDT under the same protocol as a complementary view of self-consistency. Fig. 7 reports the number of chain breaks per structure (a break is counted whenever a consecutive phosphate-phosphate distance exceeds 7.5 Å) together with the per-length break-free rate, and Fig. 8 the internal clashes per 100 heavy atoms under the MolProbity-style *d < r*_vdw,*i*_ + *r*_vdw,*j*_ − 0.6 vdW-overlap definition, the same criterion as Table 4(b). For the clash comparison, Fig. 8 additionally reports the RNAsolo training set (single-chain entries, 40-150 nt, binned at 10 nt) under the identical definition. The four figures use independent vertical scales.

**Figure 5:**
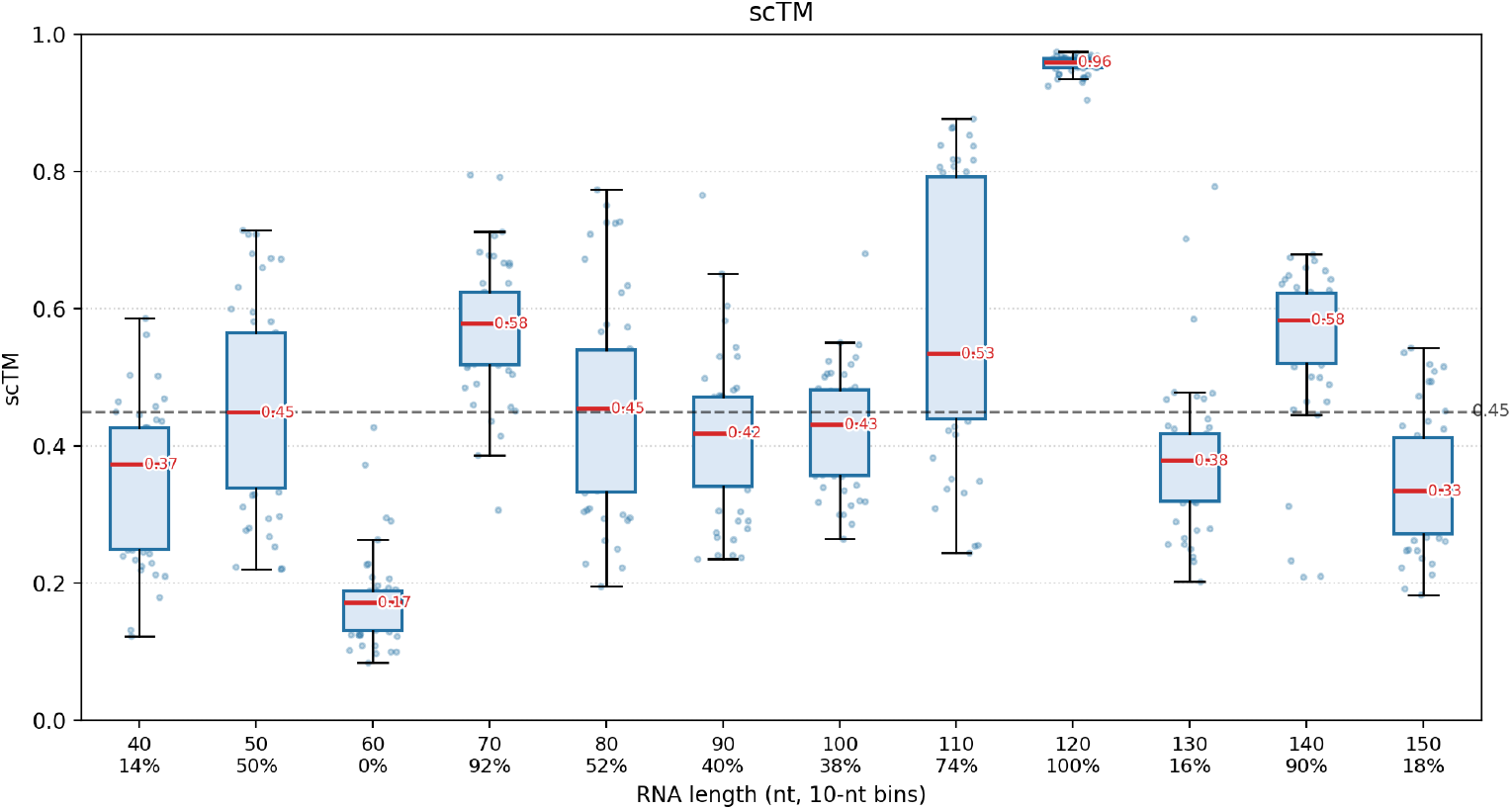
Per-length scTM distribution on the 40-150 nt grid (600 structures, 50 per length, IF protocol); the dashed line marks the scTM = 0.45 validity threshold, and the x-axis annotations give the percentage of valid structures (scTM ≥ 0.45) at each length.

**Figure 6:**
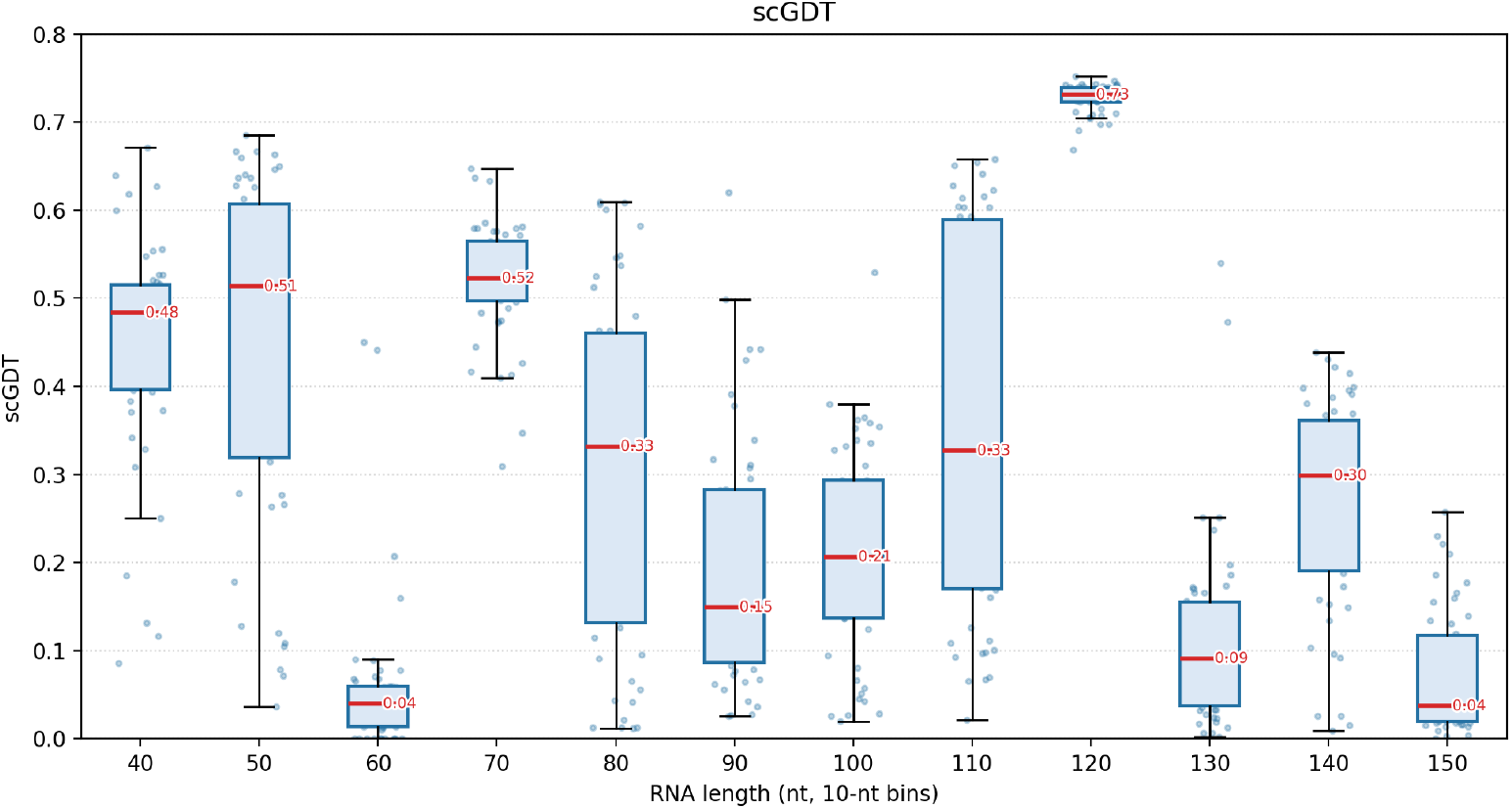
Per-length scGDT on the 40-150 nt grid under the same IF protocol (600 structures, 50 per length; each structure scored by the best of its designed sequences). Unlike scTM, no established validity threshold exists for scGDT, so no reference line is drawn.

**Figure 7:**
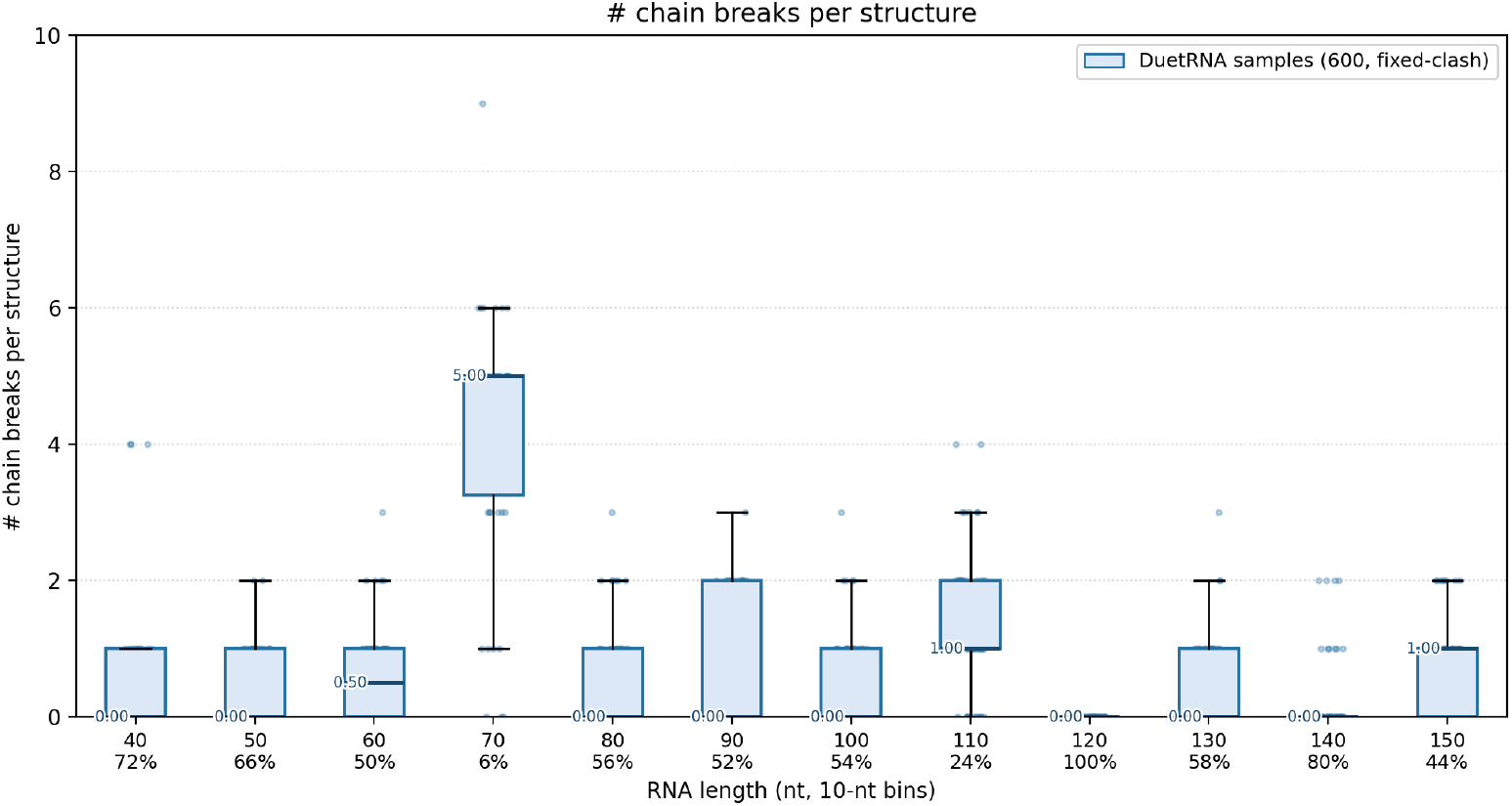
Per-length number of chain breaks per structure, where a break is counted whenever a consecutive P–P distance exceeds 7.5 Å; x-axis annotations give the percentage of break-free structures (all consecutive P-P distances ≤ 7.5 Å) at each length.

**Figure 8:**
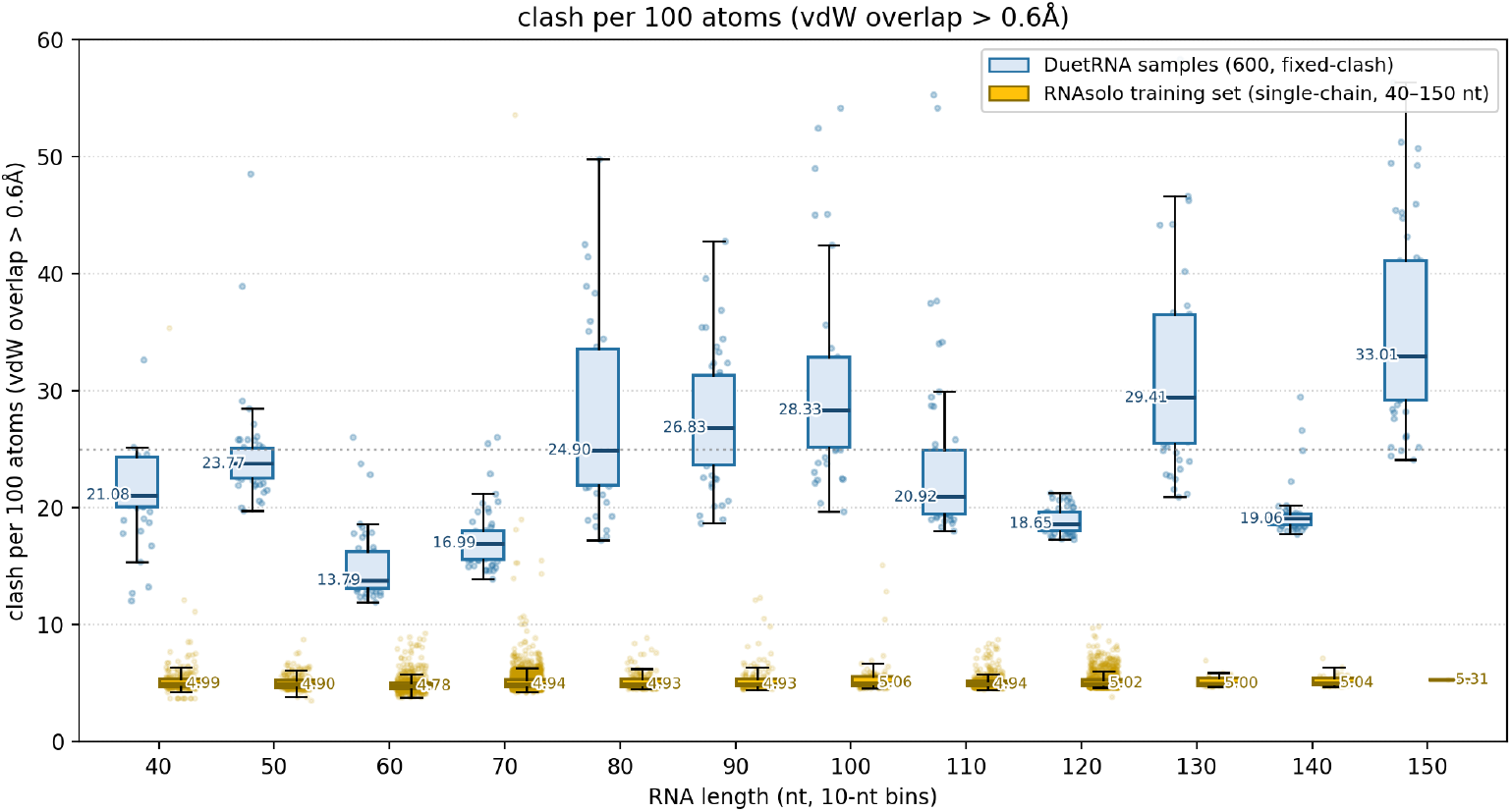
Per-length steric clashes per 100 heavy atoms under the vdW-overlap definition (*d < r*_vdw,*i*_ + *r*_vdw,*j*_ − 0.6, Bondi radii; intra-residue atom pairs and the covalent *O*3′–*P* bond excluded), the same criterion as Table 4(b). The blue boxes represent the 600 generated structures; the yellow boxes represent the RNAsolo training set restricted to single-chain entries (multi-chain concatenates excluded, since spatially overlapping chain copies inflate the all-atom clash count).

#### Self-consistency is length-dependent

The median scTM peaks at 0.959 in the 120-nt regime and drops to 0.172 at 60 nt. Recovering the target geometry through sequence alone appears harder in some length regimes. Under the single criterion scTM ≥ 0.45, 292 of the 600 structures (48.7%) are valid, with per-length validity rates ranging from 0% at 60 nt to 100% at 120 nt. The scGDT profile in Fig. 6 tracks this pattern closely (per-length medians from 0.73 at 120 nt down to 0.04 at 60 nt and 150 nt, overall median 0.28), so the length dependence of self-consistency is robust to the choice of similarity metric, although both scores are derived from the same refolded structures and therefore share common confounders.

#### Chain breaks concentrate in specific length regimes

The per-length break-free rate is non-monotonic, ranging from 6% (70 nt) to 100% (120 nt), with most regimes between 44% and 80% and an overall rate of 331*/*600 = 55.2%. The breakdown is dominated by two regimes: 70 nt, where only 6% of structures are break-free and the median structure suffers 5 breaks, and 110 nt (24% break-free, median 1 break). The fragmentation mode highlighted in Fig. 4(b) is thus localized rather than length-proportional, and the 120-nt regime used for the qualitative showcase is fully break-free.

#### Steric quality is length-uniform rather than localized

Under the 0.6 definition, the per-length median fluctuates between 13.8 (60 nt) and 33.0 (150 nt) clashes per 100 atoms with no significant trend across the twelve bins (Spearman *ρ* = 0.37, *p* ≈ 0.24), and the per-sample mean over the grid is 26.55, matching Table 4(b). Poor steric quality thus appears at all lengths, in two forms. At 40 nt, a few samples collapse into interpenetrating conformations (Fig. 4a), producing the grid’s most extreme tail (90th percentile 100). At long lengths the problem is broad rather than deep: taking the grid-wide mean of 26.55 clashes per 100 atoms as the reference level, 64% of the 130-nt and 86% of the 150-nt samples lie above it, whereas no 120-nt sample does. The gap to experimental structures is substantial at every length, consistent with the aggregate gap common to generative RNA backbones: the single-chain RNAsolo reference has per-length medians of only 4.8–5.3 clashes per 100 atoms (pooled 90th percentile 6.0), while the generated per-length medians (13.8–33.0) exceed this baseline by 2.9–6.2 × in every bin.

These per-length profiles are descriptive and complement the aggregate comparisons of §5.5. The weakest regimes are metric-specific-60 nt for self-consistency (scTM median 0.172, scGDT 0.04), 70 nt for backbone continuity (6% break-free), and the long-length bins for steric quality (with an additional collapse-driven tail at 40 nt)-and the three profiles do not co-vary with length in a common direction, so each factor follows its own regime-specific pattern that aggregate comparisons alone would mask. Finally, the training pool itself is highly imbalanced across lengths: single-chain RNAsolo entries cluster at particular length bands (1, 663 entries at 70–79 nt and 1, 026 at 120–129 nt) but are nearly absent at the long end (22 entries at 130–139 nt and a single entry at 150 nt). The degraded steric quality at 130–150 nt therefore plausibly reflects this imbalance: the model is asked to generate chains at lengths for which it has seen very few training examples.

## References

Josh Abramson, Jonas Adler, Jack Dunger, Richard Evans, Tim Green, Alexander Pritzel, Olaf Ronneberger, Lindsay Willmore, Andrew J. Ballard, Joshua Bambrick, et al. Accurate structure prediction of biomolecular interactions with AlphaFold 3. Nature, 630:493–500, 2024. doi: 10.1038/s41586-024-07487-w.

Bartosz Adamczyk, Maciej Antczak, and Marta Szachniuk. RNAsolo: a repository of cleaned PDB-derived RNA 3D structures. Bioinformatics, 38(14):3668–3670, 2022. doi: 10.1093/bioinformatics/btac386.

Rishabh Anand, Chaitanya K. Joshi, Alex Morehead, Arian Rokkum Jamasb, Charles Harris, Simon V. Mathis, Kieran Didi, Rex Ying, Bryan Hooi, and Pietro Liò. RNA-FrameFlow: Flow matching for de novo 3D RNA backbone design. Transactions on Machine Learning Research, 2025.

Minkyung Baek, Ryan McHugh, Ivan Anishchenko, Hanlun Jiang, David Baker, and Frank DiMaio. Accurate prediction of protein–nucleic acid complexes using RoseTTAFoldNA. Nature Methods, 21:117–121, 2024. doi: 10.1038/s41592-023-02086-5.

Akash Bahai, Chee Keong Kwoh, Yuguang Mu, and Yinghui Li. Systematic benchmarking of deep-learning methods for tertiary RNA structure prediction. PLOS Computational Biology, 20 (12):e1012715, 2024. doi: 10.1371/journal.pcbi.1012715.

Timothy D. Barfoot. State Estimation for Robotics. Cambridge University Press, Cambridge, UK, 2017.

Michal J. Boniecki, Grzegorz Lach, Wayne K. Dawson, Konrad Tomala, Pawel Lukasz, Tomasz Soltysinski, Kristian M. Rother, and Janusz M. Bujnicki. SimRNA: a coarse-grained method for RNA folding simulations and 3D structure prediction. Nucleic Acids Research, 44(7):e63, 2016. doi: 10.1093/nar/gkv1479.

Ricky T. Q. Chen and Yaron Lipman. Flow matching on general geometries. In International Conference on Learning Representations (ICLR), 2024.

Ricky T. Q. Chen, Yulia Rubanova, Jesse Bettencourt, and David K. Duvenaud. Neural ordinary differential equations. In Advances in Neural Information Processing Systems, volume 31, 2018.

Vincent B. Chen, W. Bryan Arendall, Jeffrey J. Headd, Daniel A. Keedy, Robert M. Immormino, Gary J. Kapral, Laura W. Murray, Jane S. Richardson, and David C. Richardson. MolProbity: all-atom structure validation for macromolecular crystallography. Acta Crystallographica Section D, 66(1):12–21, 2010. doi: 10.1107/S0907444909042073.

Mary C. Clay, Laura R. Ganser, Dawn K. Merriman, and Hashim M. Al-Hashimi. Resolving sugar puckers in RNA excited states exposes slow modes of repuckering dynamics. Nucleic Acids Research, 45(14):e134, 2017. doi: 10.1093/nar/gkx525.

José Almeida Cruz, Marc-Frédérick Blanchet, Michal Boniecki, Janusz M. Bujnicki, Shi-Jie Chen, Song Cao, Rhiju Das, Feng Ding, Nikolay V. Dokholyan, Samuel Coulbourn Flores, et al. RNA-Puzzles: A CASP-like evaluation of RNA three-dimensional structure prediction. RNA, 18(4): 610–625, 2012. doi: 10.1261/rna.031054.111.

Tulsi Ram Damase, Roman Sukhovershin, Christian Boada, Francesca Taraballi, Roderic I. Pettigrew, and John P. Cooke. The limitless future of RNA therapeutics. Frontiers in Bioengineering and Biotechnology, 9:628137, 2021. doi: 10.3389/fbioe.2021.628137.

Padideh Danaee, Mason Rouches, Michelle Wiley, Dezhong Deng, Liang Huang, and David Hendrix. bpRNA: large-scale automated annotation and analysis of RNA secondary structure. Nucleic Acids Research, 46(11):5381–5394, 2018. doi: 10.1093/nar/gky285.

Rhiju Das and David Baker. Automated de novo prediction of native-like RNA tertiary structures. Proceedings of the National Academy of Sciences, 104(37):14664–14669, 2007. doi: 10.1073/pnas.0703836104.

Andrew Favor, Riley Quijano, Elizaveta Chernova, Andrew Kubaney, Connor Weidle, Morgan A. Esler, Lilian McHugh, Ann Carr, Yang Hsia, David Juergens, et al. De novo design of RNA and nucleoprotein complexes. bioRxiv, 2025. doi: 10.1101/2025.10.01.679929.

Fabian B. Fuchs, Daniel E. Worrall, Volker Fischer, and Max Welling. SE(3)-Transformers: 3D roto-translation equivariant attention networks. In Advances in Neural Information Processing Systems, volume 33, 2020.

Laura R. Ganser, Megan L. Kelly, Daniel Herschlag, and Hashim M. Al-Hashimi. The roles of structural dynamics in the cellular functions of RNAs. Nature Reviews Molecular Cell Biology, 20 (8):474–489, 2019. doi: 10.1038/s41580-019-0136-0.

Andrew D. Garst, Andrea L. Edwards, and Robert T. Batey. Riboswitches: Structures and mechanisms. Cold Spring Harbor Perspectives in Biology, 3(6):a003533, 2011. doi: 10.1101/cshperspect.a003533.

Anke Gelbin, Bohdan Schneider, Lester Clowney, Shu-Hsin Hsieh, Wilma K. Olson, and Helen M. Berman. Geometric parameters in nucleic acids: sugar and phosphate constituents. Journal of the American Chemical Society, 118(3):519–529, 1996. doi: 10.1021/ja9528846.

Hongyu Guo, Yoshua Bengio, and Shengchao Liu. AssembleFlow: Rigid flow matching with inertial frames for molecular assembly. In International Conference on Learning Representations, 2025.

Dongran Han, Xiaodong Qi, Cameron Myhrvold, Bei Wang, Mingjie Dai, Shuoxing Jiang, Maxwell Bates, Yan Liu, Byoungkwon An, Fei Zhang, et al. Single-stranded DNA and RNA origami. Science, 358(6369):eaao2648, 2017. doi: 10.1126/science.aao2648.

Emiel Hoogeboom, Victor Garcia Satorras, Clément Vignac, and Max Welling. Equivariant diffusion for molecule generation in 3D. In Proceedings of the 39th International Conference on Machine Learning, volume 162 of Proceedings of Machine Learning Research, pages 8867–8887, 2022.

Bowen Jing, Stephan Eismann, Patricia Suriana, Raphael J. L. Townshend, and Ron O. Dror. Learning from protein structure with geometric vector perceptrons. In International Conference on Learning Representations, 2021.

Chaitanya K. Joshi, Arian Rokkum Jamasb, Ramon Viñas Torné, Charles Harris, Simon V. Mathis, Alex Morehead, Rishabh Anand, and Pietro Liò. gRNAde: Geometric deep learning for 3D RNA inverse design. In International Conference on Learning Representations, 2025.

John Jumper, Richard Evans, Alexander Pritzel, Tim Green, Michael Figurnov, Olaf Ronneberger, Kathryn Tunyasuvunakool, Russ Bates, Augustin Žídek, Anna Potapenko, et al. Highly accurate protein structure prediction with AlphaFold. Nature, 596(7873):583–589, 2021. doi: 10.1038/s41586-021-03819-2.

Yuki Kagaya, Zicong Zhang, Nabil Ibtehaz, Xiao Wang, Tsukasa Nakamura, Pranav Deep Punuru, and Daisuke Kihara. NuFold: end-to-end approach for RNA tertiary structure prediction with flexible nucleobase center representation. Nature Communications, 16:881, 2025. doi: 10.1038/s41467-025-56261-7.

Neocles B. Leontis and Eric Westhof. Geometric nomenclature and classification of RNA base pairs. RNA, 7(4):499–512, 2001. doi: 10.1017/S1355838201002515.

Haorui Li, Weitao Du, Yuqiang Li, Hongyu Guo, and Shengchao Liu. InertialAR: Autoregressive 3D molecule generation with inertial frames. In International Conference on Machine Learning (ICML), 2026a.

Xiner Li, Masatoshi Uehara, Xingyu Su, Gabriele Scalia, and Shuiwang Ji. A joint diffusion model with pre-trained priors for RNA sequence–structure co-design. In International Conference on Learning Representations, 2026b.

Yang Li, Chengxin Zhang, Chenjie Feng, Robin Pearce, P. Lydia Freddolino, and Yang Zhang. Integrating end-to-end learning with deep geometrical potentials for ab initio RNA structure prediction. Nature Communications, 14:5745, 2023. doi: 10.1038/s41467-023-41303-9.

Yang Li, Chenjie Feng, Xi Zhang, Sho Tsukiyama, Duanyu Feng, and Yang Zhang. DRfold2 is a deep learning-based tool that enables efficient and accurate RNA structure prediction. PLOS Biology, 24(2):e3003659, 2026c. doi: 10.1371/journal.pbio.3003659.

Yeqing Lin and Mohammed AlQuraishi. Generating novel, designable, and diverse protein structures by equivariantly diffusing oriented residue clouds. In Proceedings of the 40th International Conference on Machine Learning, volume 202 of Proceedings of Machine Learning Research, pages 20978–21002, 2023.

Yaron Lipman, Ricky T. Q. Chen, Heli Ben-Hamu, Maximilian Nickel, and Matt Le. Flow matching for generative modeling. In International Conference on Learning Representations (ICLR), 2023.

Xiang-Jun Lu, Harmen J. Bussemaker, and Wilma K. Olson. DSSR: an integrated software tool for dissecting the spatial structure of RNA. Nucleic Acids Research, 43(21):e142, 2015. doi: 10.1093/nar/gkv716.

Runze Ma, Zhongyue Zhang, Zichen Wang, Chenqing Hua, Jiahua Rao, Zhuomin Zhou, and Shuangjia Zheng. RiboFlow: Conditional De Novo RNA co-design via synergistic flow matching. In Advances in Neural Information Processing Systems, volume 38, pages 77810–77842, 2025. doi: 10.52202/085713-2348.

Zhichao Miao, Ryszard W. Adamiak, Maciej Antczak, Michal J. Boniecki, Janusz M. Bujnicki, Shi-Jie Chen, Clarence Yu Cheng, Yi Cheng, Fang-Chieh Chou, Rhiju Das, et al. RNA-Puzzles Round IV: 3D structure predictions of four ribozymes and two aptamers. RNA, 26(8):982–995, 2020. doi: 10.1261/rna.075341.120.

Alex Morehead, Jeffrey Ruffolo, Aadyot Bhatnagar, and Ali Madani. Towards joint sequence–structure generation of nucleic acid and protein complexes with SE(3)-discrete diffusion. arXiv preprint arXiv:2401.06151, 2023.

Laura J. W. Murray, W. Bryan Arendall, David C. Richardson, and Jane S. Richardson. RNA backbone is rotameric. Proceedings of the National Academy of Sciences, 100(24):13904–13909, 2003. doi: 10.1073/pnas.1835769100.

Zhanghan Ni, Yanjing Li, Zeju Qiu, Bernhard Schölkopf, Hongyu Guo, Weiyang Liu, and Shengchao Liu. Rigidity-aware geometric pretraining for protein design and conformational ensembles. In International Conference on Learning Representations, 2026.

Poul Nissen, Joseph A. Ippolito, Nenad Ban, Peter B. Moore, and Thomas A. Steitz. RNA tertiary interactions in the large ribosomal subunit: The A-minor motif. Proceedings of the National Academy of Sciences, 98(9):4899–4903, 2001. doi: 10.1073/pnas.081082398.

Divya Nori and Wengong Jin. RNAFlow: RNA structure & sequence design via inverse folding-based flow matching. In Proceedings of the 41st International Conference on Machine Learning, volume 235 of Proceedings of Machine Learning Research, pages 38395–38408, 2024.

Lorena G. Parlea, Blake A. Sweeney, Maryam Hosseini-Asanjan, Craig L. Zirbel, and Neocles B. Leontis. The RNA 3D Motif Atlas: Computational methods for extraction, organization and evaluation of RNA motifs. Methods, 103:99–119, 2016. doi: 10.1016/j.ymeth.2016.04.025.

Robin Pearce, Gilbert S. Omenn, and Yang Zhang. De novo RNA tertiary structure prediction at atomic resolution using geometric potentials from deep learning. bioRxiv, 2022. doi: 10.1101/2022.05.15.491755.

Mariusz Popenda, Marta Szachniuk, Maciej Antczak, Katarzyna J. Purzycka, Piotr Lukasiak, Natalia Bartol, Jacek Blazewicz, and Ryszard W. Adamiak. Automated 3D structure composition for large RNAs. Nucleic Acids Research, 40(14):e112, 2012. doi: 10.1093/nar/gks339.

Jane S. Richardson, Bohdan Schneider, Laura W. Murray, Gary J. Kapral, Robert M. Immormino, Jeffrey J. Headd, David C. Richardson, Daniela Ham, Eli Hershkovits, Loren Dean Williams, et al. RNA backbone: consensus all-angle conformers and modular string nomenclature (an RNA Ontology Consortium contribution). RNA, 14(3):465–481, 2008. doi: 10.1261/rna.657708.

Dana Rubin, Allan dos Santos Costa, Manvitha Ponnapati, and Joseph Jacobson. RiboGen: RNA sequence and structure co-generation with equivariant multiflow. arXiv preprint arXiv:2503.02058, 2025.

Wolfram Saenger. Principles of Nucleic Acid Structure. Springer-Verlag, New York, 1984. ISBN 0-387-90761-0. doi: 10.1007/978-1-4612-5190-3.

Michael Sarver, Craig L. Zirbel, Jesse Stombaugh, Ali Mokdad, and Neocles B. Leontis. FR3D: finding local and composite recurrent structural motifs in RNA 3D structures. Journal of Mathematical Biology, 56(1–2):215–252, 2008. doi: 10.1007/s00285-007-0110-x.

Victor Garcia Satorras, Emiel Hoogeboom, and Max Welling. E(n) equivariant graph neural networks. In Proceedings of the 38th International Conference on Machine Learning, volume 139 of Proceedings of Machine Learning Research, pages 9323–9332, 2021.

Kristof T. Schütt, Oliver T. Unke, and Michael Gastegger. Equivariant message passing for the prediction of tensorial properties and molecular spectra. In Proceedings of the 38th International Conference on Machine Learning, volume 139 of Proceedings of Machine Learning Research, pages 9377–9388, 2021.

Tao Shen, Zhihang Hu, Zhangzhi Peng, Jiayang Chen, Peng Xiong, Liang Hong, Liangzhen Zheng, Yixuan Wang, Irwin King, Sheng Wang, Siqi Sun, and Yu Li. E2Efold-3D: End-to-end deep learning method for accurate de novo RNA 3D structure prediction, 2022.

Tao Shen, Zhihang Hu, Siqi Sun, D. Liu, Felix Wong, Jiuming Wang, Jiayang Chen, Yixuan Wang, Liang Hong, Jin Xiao, Liangzhen Zheng, Tejas Krishnamoorthi, Irwin King, Sheng Wang, Peng Yin, James J. Collins, and Yu Li. Accurate RNA 3D structure prediction using a language model-based deep learning approach. Nature Methods, 21:2287–2298, 2024. doi: 10.1038/s41592-024-02487-0.

Marcell Szikszai, Marcin Magnus, Siddhant Sanghi, Sachin Kadyan, Nazim Bouatta, and Elena Rivas. RNA3DB: A structurally-dissimilar dataset split for training and benchmarking deep learning models for RNA structure prediction. Journal of Molecular Biology, 436(17):168552, 2024. doi: 10.1016/j.jmb.2024.168552.

Sumit Tarafder and Debswapna Bhattacharya. RNAbpFlow: base pair-augmented SE(3) flow matching for conditional RNA 3D structure generation. Nature Methods, 23:1349–1358, 2026. doi: 10.1038/s41592-026-03128-4.

Raphael J. L. Townshend, Stephan Eismann, Andrew M. Watkins, Ramya Rangan, Masha Karelina, Rhiju Das, and Ron O. Dror. Geometric deep learning of RNA structure. Science, 373(6558): 1047–1051, 2021. doi: 10.1126/science.abe5650.

Quentin Vicens and Jeffrey S. Kieft. Thoughts on how to think (and talk) about RNA structure. Proceedings of the National Academy of Sciences, 119(17):e2112677119, 2022. doi: 10.1073/pnas.2112677119.

Jun Wang, Jian Wang, Yanzhao Huang, and Yi Xiao. 3dRNA v2.0: An updated web server for RNA 3D structure prediction. International Journal of Molecular Sciences, 20(17):4116, 2019. doi: 10.3390/ijms20174116.

Wenkai Wang, Chenjie Feng, Renmin Han, Ziyi Wang, Lisha Ye, Zongyang Du, Hong Wei, Fa Zhang, Zhenling Peng, and Jianyi Yang. trRosettaRNA: automated prediction of RNA 3D structure with transformer network. Nature Communications, 14:7266, 2023. doi: 10.1038/s41467-023-42528-4.

Andrew Martin Watkins, Ramya Rangan, and Rhiju Das. FARFAR2: Improved de novo Rosetta prediction of complex global RNA folds. Structure, 28(8):963–976.e6, 2020. doi: 10.1016/j.str.2020.05.011.

Joseph L. Watson, David Juergens, Nathaniel R. Bennett, Brian L. Trippe, Jason Yim, Helen E. Eisenach, Woody Ahern, Andrew J. Borst, Robert J. Ragotte, Lukas F. Milles, et al. De novo design of protein structure and function with RFdiffusion. Nature, 620:1089–1100, 2023. doi: 10.1038/s41586-023-06415-8.

Christopher J. Williams, Jeffrey J. Headd, Nigel W. Moriarty, Michael G. Prisant, Lizbeth L. Videau, Lindsay N. Deis, Vishal Verma, Daniel A. Keedy, Bradley J. Hintze, Vincent B. Chen, et al. MolProbity: More and better reference data for improved all-atom structure validation. Protein Science, 27(1):293–315, 2018. doi: 10.1002/pro.3330.

Jeremy Wohlwend, Gabriele Corso, Saro Passaro, Noah Getz, Mateo Reveiz, Ken Leidal, Wojtek Swiderski, Liam Atkinson, Tally Portnoi, Itamar Chinn, Jacob Silterra, Tommi Jaakkola, and Regina Barzilay. Boltz-1: Democratizing biomolecular interaction modeling. bioRxiv, 2024. doi: 10.1101/2024.11.19.624167. Preprint.

Joseph D. Yesselman, Daniel Eiler, Erik D. Carlson, Michael R. Gotrik, Anne E. d’Aquino, Alexandra N. Ooms, Wipapat Kladwang, Paul D. Carlson, Xuesong Shi, David A. Costantino, et al. Computational design of three-dimensional RNA structure and function. Nature Nanotechnology, 14(9):866–873, 2019. doi: 10.1038/s41565-019-0517-8.

Jason Yim, Andrew Campbell, Andrew Y. K. Foong, Michael Gastegger, José Jiménez-Luna, Sarah Lewis, Victor Garcia Satorras, Bastiaan S. Veeling, Regina Barzilay, Tommi Jaakkola, et al. Fast protein backbone generation with SE(3) flow matching. arXiv preprint arXiv:2310.05297, 2023a.

Jason Yim, Brian L. Trippe, Valentin De Bortoli, Emile Mathieu, Arnaud Doucet, Regina Barzilay, and Tommi Jaakkola. SE(3) diffusion model with application to protein backbone generation. In Proceedings of the 40th International Conference on Machine Learning, volume 202 of Proceedings of Machine Learning Research, pages 40001–40039, 2023b.

Chengxin Zhang, Morgan Shine, Anna Marie Pyle, and Yang Zhang. US-align: universal structure alignments of proteins, nucleic acids, and macromolecular complexes. Nature Methods, 19(9): 1109–1115, 2022. doi: 10.1038/s41592-022-01585-1.

Yize Zhou, Haorui Li, and Shengchao Liu. A resolution-agnostic geometric transformer for chromosome modeling using inertial frame. In International Conference on Learning Representations, 2026.

